# Cell type-specific biotin labeling *in vivo* resolves regional neuronal proteomic differences in mouse brain

**DOI:** 10.1101/2021.08.03.454921

**Authors:** Sruti Rayaprolu, Sara Bitarafan, Ranjita Betarbet, Sydney Sunna, Lihong Cheng, Hailian Xiao, Pritha Bagchi, Duc M. Duong, Ruth Nelson, Annie M. Goettemoeller, Viktor János Oláh, Matt Rowan, Allan I. Levey, Levi B. Wood, Nicholas T. Seyfried, Srikant Rangaraju

## Abstract

Isolation and proteomic profiling of brain cell types, particularly neurons, pose several technical challenges which limit our ability to resolve distinct cellular phenotypes in neurological diseases. Therefore, we generated a novel mouse line that enables cell type-specific expression of a biotin ligase, TurboID, via Cre-lox strategy for *in vivo* proximity-dependent biotinylation of proteins. Using adenoviral-based and transgenic approaches, we show striking protein biotinylation in neuronal cell bodies and axons throughout the mouse brain. We quantified more than 2,000 neuron-derived proteins following enrichment that mapped to numerous subcellular compartments. Synaptic, transmembrane transporters, ion channel subunits, and disease-relevant druggable targets were among the most significantly enriched proteins. Remarkably, we resolved brain region-specific proteomic profiles of Camk2a neurons with distinct functional molecular signatures and disease associations that may underlie regional neuronal vulnerability. Leveraging the neuronal specificity of this *in vivo* biotinylation strategy, we used an antibody-based approach to uncover regionally unique patterns of neuron-derived signaling phospho-proteins and cytokines, particularly in the cortex and cerebellum. Our work provides a proteomic framework to investigate cell type-specific mechanisms driving physiological and pathological states of the brain as well as complex tissues beyond the brain.

## INTRODUCTION

The brain is comprised of distinct cell types including neurons, glia (astrocytes, oligodendrocytes, microglia), and vascular cells (endothelial cells and pericytes). Brain cell types are immensely complex and have nuanced changes in their molecular composition (DNA, mRNA, protein) during physiological processes in development, aging, and in response to pathological insults^1^. Rapid advances in bulk and single cell transcriptomic profiling of healthy and pathogenic states of mouse and human brain have provided numerous new insights into diverse molecular signatures adopted by brain cell types^2^. Specifically, they have highlighted several causal neuronal and glial cellular responses in neurological diseases; however, transcriptomic findings only modestly correlate with protein-level (proteomic) changes^3–6^. Proteins also undergo post-translational modifications that impact their function and expression profiles quantitatively, temporally, and spatially; a feature not reflected by transcript abundance^3,5,6^. Therefore, proteomic characterization of distinct brain cell types can provide important insights into cellular mechanisms of development, aging and neuropathology.

Proteomic profiling of brain cell types requires isolation of intact cells from fresh unfrozen brain, a pre-requisite that is not necessary for single-nuclear transcriptomic studies of intact nuclei from frozen brain^7,8^. There are numerous methods by which relatively pure, live cells of interest can be isolated with minimal contamination and high sensitivity. For instance, magnetic activated cell sorting (MACS) and fluorescence activated cell sorting (FACS) have been used to isolate one or more brain cell type for mass spectrometry (MS)-based proteomics^9,10^. However, these methods are not without limitations, including variable cell yield, low protein yields, cellular and acellular contamination, sampling biases, and artefactual cellular activation, which significantly impact the proteomic profiles of these cell types^9^. Among brain cell types, live neurons from fresh brain tissue are particularly difficult to isolate, inherently limiting our ability to understand neuron-specific proteomic changes occurring *in vivo*.

Recently, *in vivo* cell type-specific transcriptomic labeling approaches (e.g., RiboTag) were developed to quantify transcripts without the need for cell isolation^11–13^. Analogous to these approaches, *in vivo* proteomic labeling has been achieved by bio-orthogonal non-canonical amino acid tagging (BONCAT) where a Cre-lox genetic strategy leads to conditional expression of the mutant (L274G) methionyl tRNA synthetase (MetRS*) in a specific cell type in the mouse brain^14^. MetRS* incorporates a methionine analog, azidonorleucine (Anl), into newly synthesized proteins in place of methionine in the desired cell type^14–17^. Anl-tagged proteins can then be enriched using azide-alkyne “Click” chemistry followed by MS quantitation^17^. Thus far, *in vivo* BONCAT has been successfully applied to characterizing the nascent neuronal proteome and is well-suited to study turnover of newly synthesized proteins^14,15^.

Proximity-dependent protein labeling approaches using highly efficient and promiscuous biotin ligases, such as TurboID, represent an alternative approach for global proteomic labeling within a cell^18^. TurboID is a highly efficient biotin ligase that can label proteins in a 9-10nm radius within 10 min and at low temperatures^19^. This allows for the investigation of dynamic biological processes with greater proteomic breadth. *In vivo* applications of TurboID have been limited to investigating protein-protein interactions by fusing TurboID to the protein of interest or obtaining peripheral cell type-specific secretome profiles^19^. In the context of the brain, the proteomic profiles of astrocyte-synapse interactions were characterized in the mouse brain by introducing TurboID via adeno-associated viruses (AAV)^20^, but brain cell type-specific proteomics using TurboID has been limited by lack of a suitable mouse model.

Here, we developed and validated a *Rosa26^TurboID^* mouse line for TurboID expression via Cre-lox genetic approaches to induce TurboID expression in neurons. We term this approach <u>c</u>ell type-specific *in vivo* <u>b</u>iotinylation <u>o</u>f <u>p</u>roteins or CIBOP. We first used an AAV mediated delivery of Cre recombinase under a synapsin promoter for pan-neuronal labeling of proteins in the hippocampus. Our second application involved a genetic approach by crossing the *Rosa26^TurboID^* mouse with Camk2a-Cre^Ert^^2^ mice to label Camk2a neurons in the entire brain in an inducible manner. AAV-CIBOP and Tg-CIBOP resulted in robust biotinylation of proteins specifically within neuronal cell bodies and axons in the mouse brain without histological or electrophysiological abnormalities. We successfully enriched these biotinylated proteins from total brain homogenates and captured the neuronal proteome (>2,000 proteins) by MS. Furthermore, we were able to resolve unique proteomic signatures of Camk2a neurons that are region-specific and indicative of distinct cellular functions and disease-vulnerability. Lastly, we complemented our untargeted MS approach with a targeted measurement of biotinylated phospho-proteins from key cellular signaling pathways (MAPK and Akt/mTOR) as well as biotinylated cytokines that are often below MS detection limits. This allowed us to quantify neuron-derived signaling phosphoproteins and cytokines with novel regional expression patterns in the brain. The *Rosa26^TurboID^* mouse, our validated approaches for cell type-specific *in vivo* proteomic labeling, and optimized workflow for proteomic characterization represent a highly promising approach for global cellular proteomics of desired cell types in their native state *in vivo* without the need for cell type isolation.

## RESULTS

### Hippocampal pan-neuronal proteomics using an adeno-associated viral (AAV) strategy

In a cohort of 3-month-old *Rosa26^TurboID/wt^* mice (**Fig 1a**) and wild-type (WT) littermate controls, we stereotaxically injected AAV9 carrying the Cre recombinase gene under the human synapsin (hSyn) promoter (AAV9-hSyn-Cre) into bilateral hippocampii for pan-neuronal TurboID expression (**Fig 1b**). Un-injected WT mice also served as controls (**Fig 1b**). Following 4 weeks, mice received biotin supplementation in water (37.5mg/L)^21^ for 2 weeks with no observable adverse effects (e.g., weight and locomotor activity). Immunofluorescence imaging of hippocampal regions confirmed neuronal (NeuN) biotinylation (detected by streptavidin Alexa 488) in cell bodies, axons, and synapses in majority of hippocampal CA2/3 neurons (**Fig 1c**). Western blot analysis of forebrain lysates from mice confirmed TurboID expression (based on V5 detection) and robust biotinylation of proteins in the *Rosa26^TurboID/wt^*/hSyn mice as compared to minimal endogenous biotinylation observed in control mice (**Fig 1d**).

**Figure 1.**
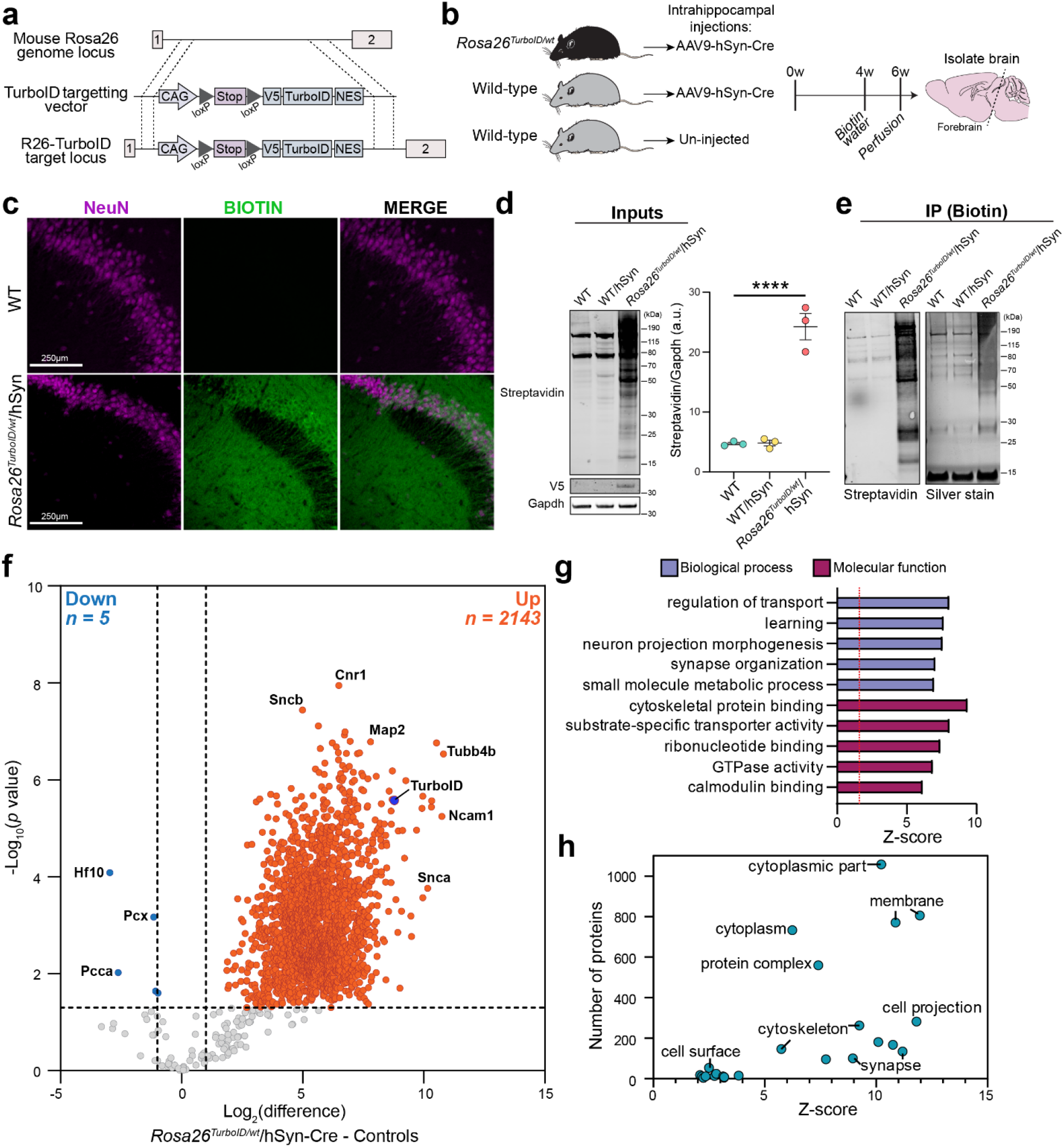
Development and validation of a novel *Rosa26^TurboID/wt^* mouse line for cell type-specific proteomics. **a** Genetic strategy for targeting TurboID (V5-TurboID-NES) to the Rosa26 locus under the CAG promoter. **b** Schematic of AAV studies to direct Cre recombinase expression in neurons. Bilateral stereotaxic hippocampal injections were performed (*n* = 3 WT mice with no injections, *n* = 3 WT mice received AAV9-hSyn-Cre, *n* = 3 *Rosa26^TurboID/wt^* mice received AAV9-hSyn-Cre), followed by 4 weeks for transduction, another 2 weeks of biotin water supplementation, and then euthanasia. **c** Representative immunofluorescence images of CA2/3 of the hippocampus from control and *Rosa26^TurboID/wt^*/hSyn showing biotinylation (green: streptavidin Alexa488) in relation to neuronal nuclei (magenta: NeuN). **d** Western blot of brain lysates from three experimental groups (1 representative animal shown) probed with streptavidin fluorophore, anti-V5, and anti-Gapdh antibodies. *Rosa26^TurboID/wt^*/hSyn brain showed biotinylated proteins of different molecular weights as compared to few endogenously biotinylated proteins in the two control groups. Right: Densitometry confirming significant increase in biotinylation signal in *Rosa26^TurboID/wt^*/hSyn brains (one-way ANOVA, \*\*\*\**p*≤0.0001). **e** Western blot (left) and silver stain (right) of enriched biotinylated proteins after strept avidin-pulldown and release of biotinylated proteins from 10% of streptavidin beads. As compared to minimal protein enriched in the two control groups, several biotinylated proteins were enriched from *Rosa26^TurboID/wt^*/hSyn brain. **f** Volcano plots showing differentially enriched proteins comparing *Rosa26^TurboID/wt^*/hSyn (*n* = 3) and control mice (*n* = 3 WT and *n* = 3 WT/hSyn). Orange symbols (*p* 0.05 and 2-fold change) represent biotinylated proteins enriched in the *Rosa26^TurboID/wt^*/hSyn brain and examples of neuron-specific proteins are highlighted, in addition to TurboID. Blue symbols represent endogenously biotinylated carboxylases enriched in the control brains. For group wise comparisons, see Supplemental Figure 1. **g** Results from GSEA of ≥ 2-fold biotinylated neuronal proteins (orange symbols from panel f), as compared to reference list^5^ (mouse brain: *n* = 7736) showed enrichment of neuronal and synaptic proteins confirming neuron-specific labeling. **h** Graphical representation of the number of proteins within various cellular compartments determined from GSEA. For related MS data and additional analyses, see Supplemental Tables 1, 3, 7, & 8.

Biotinylated proteins were enriched from forebrain lysates using streptavidin beads. Enrichment was confirmed by eluting a small fraction of bound proteins for confirmatory Western blot and silver stain studies, which showed maximum biotinylated protein yield in the *Rosa26^TurboID/wt^*/hSyn mice (**Fig 1e**). After label-free quantitation via mass spectrometry (LFQ-MS) of enriched biotinylated proteins, we quantified 2,694 unique proteins across all samples. After imputing missing values, 2,307 proteins remained in our analyses. We first compared *Rosa26^TurboID/wt^*/hSyn mice with all control mice and identified 2,143 proteins with a significant (*p* ≤ 0.05) ≥ 2-fold enrichment and 7 proteins with ≥ 2-fold enrichment in the control proteome (**Fig 1f, Supp Table 1**). The seven proteins enriched in the control samples included endogenously biotinylated carboxylases and keratins, indicative of competition between TurboID-mediated and endogenously biotinylated proteins for streptavidin binding. By including phosphorylation of Ser/Thr/Tyr residues in the searches, we identified 147 phospho-peptides from 110 unique proteins of which 55 showed at least 2-fold higher levels in the labeled neuronal proteome (e.g., Map2, Ncam1, Mapt, Map1b, Amph, **Supp Table 2**), highlighting the ability of CIBOP to capture abundant phosphorylated proteoforms even without phosphopeptide enrichment.

Gene set enrichment analyses (GSEA) of the 2,143 enriched biotinylated proteins revealed over-representation of neuronal and synaptic proteins involved in neuron projection and synapse organization, confirming neuron-specificity of labeling (**Fig 1g-h, Supp Table 3**). GSEA also showed labeling of proteins involved in a diverse set of molecular functions such as cytoskeletal protein binding, substrate-specific transporter activity, and calmodulin binding. Specifically, we observed biotinylation of 54 cell surface proteins, 63 transmembrane transporters, and 45 ion channel subunits (7 calcium, 9 glutamate, 6 GABA, 13 potassium, 1 sodium, 4 anionic) (**Supp Table 4**). The biotinylated proteome was enriched in cytoplasmic, membrane, synaptic, and cytoskeletal proteins, which are representative of the whole cell proteome rather than a bias to a specific subcellular compartment (**Fig 1h, Supp Table 3**). In comparison to bulk brain proteomic data from the AAV cohort, our pan-neuronal enriched proteome identified 354 neuron-derived proteins (**Supp Table 5**) that were not previously quantified at the whole brain level (**Supp Table 6**). GSEA of these neuron-derived proteins showed that they were predominantly membrane proteins involved in a variety of molecular functions such as serotonin receptor, potassium channel, glutamate receptor, and protein kinase activity (**Supp Table 7**). Within the 354 proteins, we identified 33 druggable targets of disease relevance (**Supp Table 5**). Druggable protein targets included solute transporters, GPCRs (e.g., GRM1), lipid metabolic proteins (e.g., GBA2), and signaling proteins including AKT1 which is a key member of the IGF1/PI3K/AKT1/mTOR axis that is relevant to synaptic functioning and memory^22^. We also performed all pairwise comparisons between labeled and control mice and obtained nearly identical GSEA results as above (**Supp Fig 1d-e and Supp Tables 1, 8, & 9**).

In sum, AAV-CIBOP resulted in robust pan-neuronal labeling of proteins in the hippocampus by TurboID *in vivo*. Proteomic analysis of biotinylated proteins confirmed neuronal enrichment and representation of proteins within a diverse number of molecular functions from various cellular compartments, including numerous synaptic proteins, transmembrane proteins, and several druggable targets which are otherwise challenging to sample in the native state of neurons in adult mouse brain.

### Camk2a-neuronal biotinylation in adult mouse brain using a transgenic strategy

We next employed a transgenic approach to express TurboID within Camk2a neurons by breeding *Rosa26^TurboID/wt^* mice with Camk2a-Cre^Ert2^ and inducing Cre recombinase expression by administering tamoxifen intraperitoneally. Camk2a (Ca^2+^/calmodulin-activated protein kinase 2A) is an abundant serine-threonine kinase highly expressed by excitatory neurons particularly in the synapse, where it regulates synaptic transmission, excitability, and long-term potentiation. Camk2a was chosen based on extensive validation, specificity, and non-leakiness of available Camk2a-Cre^Ert2^ driver lines and the well-characterized expression patterns of Camk2a across brain regions^23,24^. *Rosa26^TurboID/wt^*/Camk2a and littermate controls received tamoxifen at 6 weeks of age followed by biotin supplementation for 2 weeks (**Fig 2a**). There was no associated phenotypic changes or observable adverse effects during biotin supplementation (e.g., weight and locomotor activity). Western blot analysis of lysates from different brain regions (cortex, hippocampus, striatum/thalamus, pons/medulla, and cerebellum) confirmed robust biotinylation of proteins in the *Rosa26^TurboID/wt^*/Camk2a mice as compared to minimal endogenous biotinylation observed in control mice (**Fig 2b**). Qualitatively, the highest level of labeling was observed in the cortex, hippocampus, and striatum/thalamus regions as compared to cerebellum and pons/medulla. We also confirmed TurboID protein expression via detection of V5 in *Rosa26^TurboID/wt^*/Camk2a brain regions only, which followed a similar pattern to level of biotinylation (**Fig 2b**).

**Figure 2.**
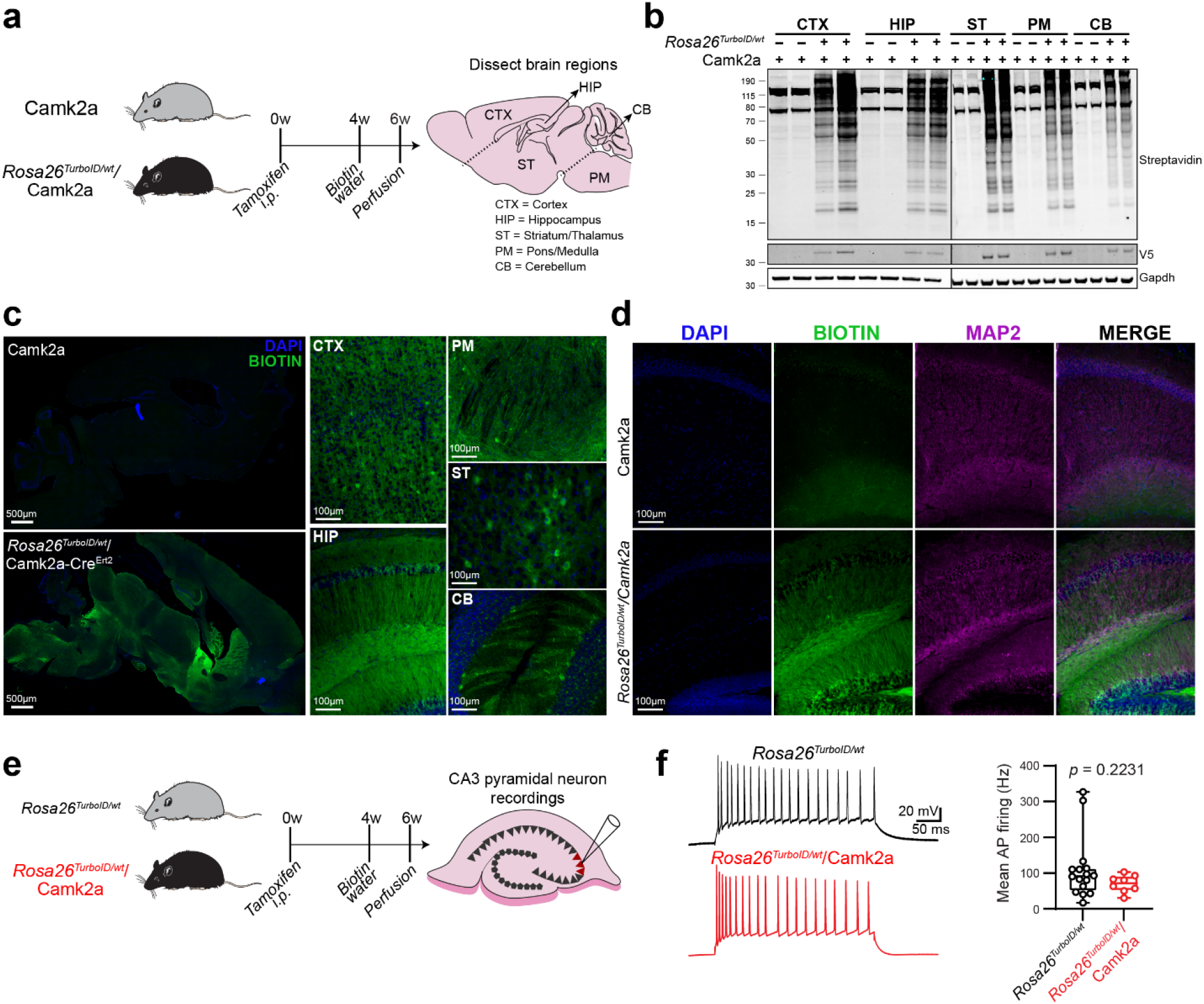
Successful biotin labeling of Camk2a neurons in adult mouse brain. **a** Schematic displaying the study design for achieving Camk2a neuron-specific proteomic biotinylation. Heterozygous Camk2a-Cre^Ert2^ (littermate controls) and *Rosa26^TurboID/wt^*/Camk2a-Cre^Ert2^ mice received tamoxifen intraperitoneally (i.p.) once a day for 5 days. After waiting 4 weeks for sufficient recombination and washout of tamoxifen effect, mice were supplemented with biotin water (37.5 mg/L) for 2 weeks. Brain was dissected into the following regions: cortex (CTX), hippocampus (HIP), striatal region which included the thalamus (ST), brain stem comprised on pons/medulla (PM), and cerebellum (CB). **b** Western blots from brain lysates, probed for biotin (streptavidin fluorophore), V5 (to detect V5-TurboID-NES), and Gapdh are shown. *Rosa26^TurboID/wt^*/Camk2a brain regions showed biotinylated proteins of different molecular weights as compared to few endogenously biotinylated proteins in the control brain regions. **c** Representative immunofluorescence images showing biotinylation (green: streptavidin Alexa488) across different brain regions. Tiled image of the entire hemisphere (sagittal section) from control and labeled mice, and higher-magnification images from individual regions of labeled mice are shown. Nuclei were labeled with DAPI (blue). **d** Representative immunofluorescence images showing overlap between biotinylation (green: streptavidin Alexa488) and Map2 protein expression (magenta) in neurons and axons in the hippocampus region from control and labeled mice. Nuclei were labeled with DAPI (blue). **e** Experimental outline for hippocampal slice electrophysiology from non-labeled *Rosa26^TurboID/wt^* (littermate control) and labeled *Rosa26^TurboID/wt^*/Camk2a mice (*n* = 2 mice /group). **f** Electrophysiological recordings from CA3c pyramidal neurons of the hippocampus showing similar firing pattern in controls and labeled *Rosa26^TurboID/wt^*/Camk2a mice. Each data point represents a single neuron. Pooled analysis from *n* = 17 non-labeled control and *n* = 8 labeled neurons is shown on the right (*p* value represents unpaired t-test). For associated immunofluorescence images for confirmation of efficient biotin labeling, and additional electrophysiological studies performed, see Supplemental Figure 4.

Immunofluorescence imaging of the whole brain displayed wide-spread biotinylation in *Rosa26^TurboID/wt^*/Camk2a brains compared to control brains (**Fig 2c**). Furthermore, biotinylation was observed predominantly within cell bodies and axonal projections in all brain regions sampled for proteomic analysis (**Fig 2c**). Map2 and streptavidin co-immunofluorescence confirmed neuronal labeling throughout the hippocampus (**Fig 2d**), as well as the cortex (**Supp Fig 2a**), pons/medulla (**Supp Fig 2b**), striatum/thalamus (**Supp Fig 2c**), and cerebellum (**Supp Fig 2d**). Importantly, biotinylation was not observed in astrocytes (**Supp Fig 3a**) or microglia (**Supp Fig 3b**), with no evidence of reactive gliosis, in *Rosa26^TurboID/wt^*/Camk2a mice compared to control mice (**Supp Fig 3**).

To determine whether TurboID expression and protein biotinylation impact Camk2a neuron activity, we performed electrophysiological studies on 3-month-old *Rosa26^TurboID/wt^*/Camk2a mice and littermate controls that received tamoxifen and biotin supplementation (**Fig 2e**). Somatic whole-cell recordings from pyramidal neurons in the CA3c region of the hippocampus, a region which displayed robust biotinylation (**Supp Fig 4a**), showed no significant differences in mean action potential firing between control and *Rosa26^TurboID/wt^*/Camk2a mice (**Fig 2f**). Furthermore, we did not observe significant differences in various passive and active parameters measured (**Supp Fig 4b**). In summary, these findings validate the Tg-CIBOP approach and confirm lack of any electrophysiological perturbations in Camk2a neurons despite robust proteomic biotinylation as well as the lack of phenotypic changes in the mouse.

### Proteomic analysis reveals unique Camk2a neuron brain region signatures

After confirming biotinylation of proteins by Western blot and immunofluorescence imaging in *Rosa26^TurboID/wt^*/Camk2a brains, we enriched for biotinylated proteins from cortex, hippocampus, striatum/thalamus, pons/medulla, and cerebellum using streptavidin beads, followed by LFQ-MS (**Fig 3a**). We quantified 2,862 unique proteins across all samples and after imputing missing values, 2,096 proteins remained in our analyses. Differential expression analysis of proteomic data, after normalizing for TurboID levels across brain regions, from all brain regions comparing *Rosa26^TurboID/wt^*/Camk2a and controls resulted in 1,245 proteins with ≥ 2-fold enrichment and 4 proteins with ≥ 2-fold enrichment in the control brains (**Supp Fig 5a, Supp Table 10**). GSEA of 1,245 enriched proteins identified proteins involved in biological processes such as transport, cytoskeletal organization, and cellular membrane organization as well as proteins involved in molecular functions such as cytoskeletal protein binding, glutamate receptor activity, and clathrin binding (**Supp Fig 5b, Supp Table 11**). Proteins within the cytoplasm, membrane, neuronal cell body, synapse, and other cellular components were significantly enriched in the *Rosa26^TurboID/wt^*/Camk2a proteome (**Supp Fig 5c**). In comparison to a reference bulk brain proteomic dataset from adult mouse brain^5^, our Camk2a neuronal proteome identified 114 neuron-derived proteins that were not previously quantified at the whole brain level (**Supp Table 12**).

**Figure 3.**
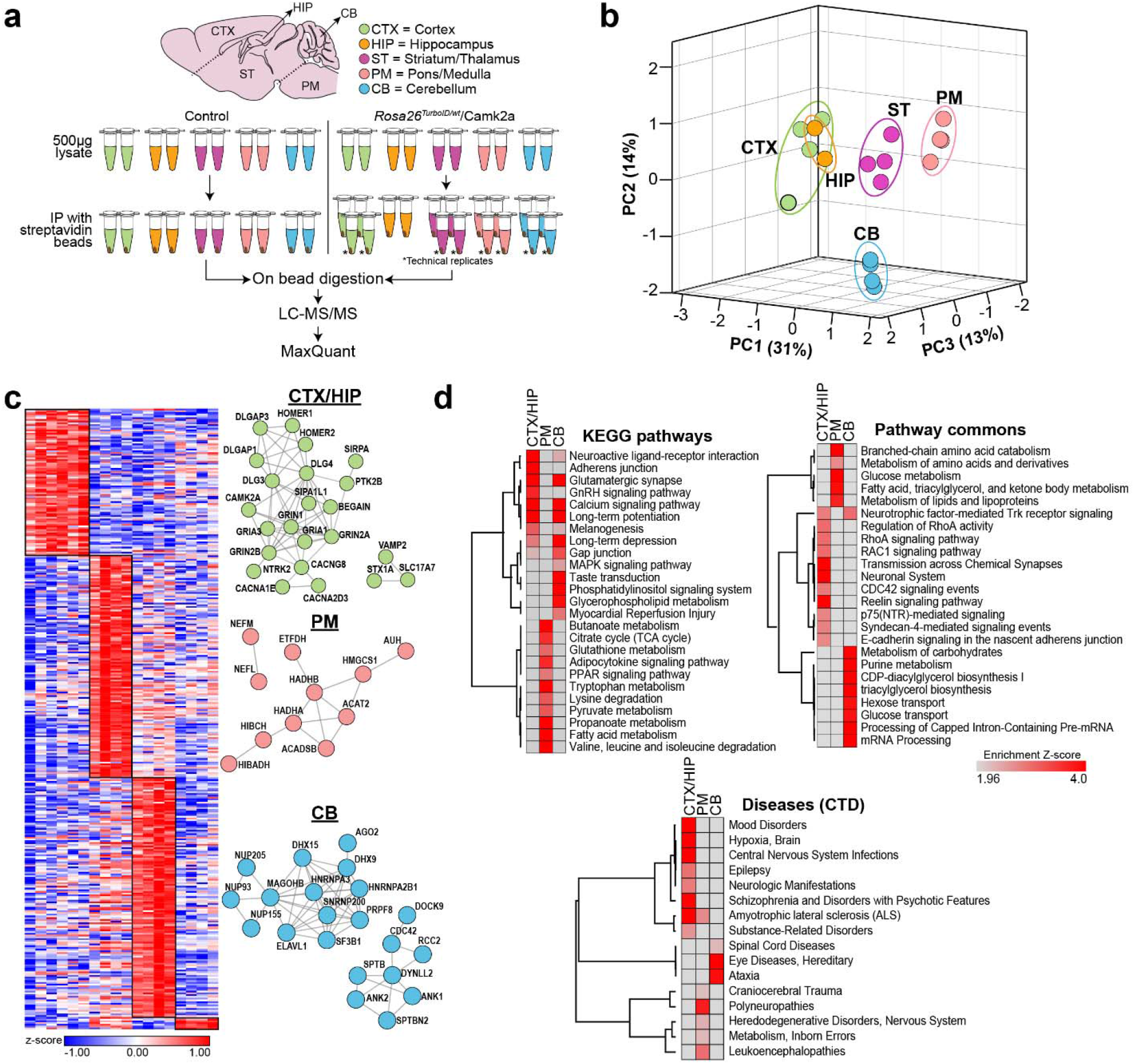
Camk2a neurons exhibit region-specific proteomic differences in adult mouse brain. **a** Experimental outline for LFQ-MS studies performed on biotinylated proteins enriched using streptavidin beads from *Rosa26^TurboID/wt^*/Camk2a-Cre^Ert2^ and littermate control (Camk2a-Cre^Ert2^) mice (*n* = 2 mice per experimental group, 5 regions per mouse). Asterisk (*) indicates technical replicates for each brain region from *Rosa26^TurboID/wt^*/Camk2a-Cre^Ert2^ mice. **b** Principal Component Analysis (PCA) of MS data after normalization to TurboID abundance in each sample/brain region. PCA identified distinct clusters based on region except for hippocampal and cortical regions clustering together. Three PCs explained 30%, 14% and 13% of variance, respectively. **c** Clustering representation of protein abundance data of core groups of proteins most highly expressed in specific brain regions with at least 4-fold higher levels in a specific region compared to all other regions (*p* ≤ 0.05). STRING analysis identified networks of known direct (protein-protein) and indirect (functional) interactions within core regional protein signatures. **d** Heatmap representation, based on enrichment Z-scores, of KEGG pathways, Pathway Commons, and diseases from the Comparative Toxicogenomics Database (CTD) enriched in core regional proteins.

Principal component analysis (PCA) of *Rosa26^TurboID/wt^*/Camk2a biotinylated protein abundance data showed that nearly 60% of variance was explained by 3 principal components (PCs), of which PC1 explained 30% while PC2 explained 14% and PC3 explained 13% of variance (**Fig 3b**). Notably, PCA identified four distinct brain regional proteomic clusters: cortex and hippocampus, striatum/thalamus, pons/medulla, and cerebellum (**Fig 3b**). We identified core groups of highly expressed region-specific proteomic signatures of Camk2a neurons (≥ 4-fold change and *p* ≤ 0.05, **Fig 3c**, **Supp Table 13**). The cortex/hippocampus contained 83 signature proteins while cerebellum contained 138, pons/medualla contained 128, and striatum/thalamus contained 6. We also identified networks of known direct (protein-protein) and indirect (functional) interactions (STRING) within core regional protein signatures (**Fig 3c**).

GSEA of core protein signatures revealed distinct molecular and biological features of Camk2a neurons (**Supp Tables 14 & 15**). The proteomic signature of cortical and hippocampal neurons was indicative of glutamatergic synaptic transmission, calcium signaling, synaptic plasticity, and more specifically, E-cadherin and syndecan-4-mediated signaling and RhoA activity (**Fig 3d**). Cortical and hippocampal neurons signature proteins were enriched in genes associated with epilepsy, mood and psychotic disorders, substance abuse, and hypoxic brain injury (**Fig 3d**). The cerebellar Camk2a proteomic signature was indicative of increased MAPK signaling and mRNA processing (splicing, snRNP assembly) as well as genes related to ataxia, spinal cord disease, and hereditary eye diseases (e.g., retinitis pigmentosa) (**Fig 3d**). Unlike cortex and cerebellar neurons, Camk2a neurons from the pons/medulla showed increased levels of proteins involved in amino acid, fatty acid, and carbohydrate metabolism (**Fig 3d**), supportive of elevated metabolic activity in these neurons and axons. Consistent with the abundant metabolic protein signature of pons/medullary Camk2a neurons and the axonal predominance in this region, this regional proteomic signature was enriched in gene symbols related to inborn errors of metabolism, leukoencephalopathies, and polyneuropathies (**Fig 3d**). Since ion channels determine the electrophysiological properties of neurons, we also identified distinct ion channel subunits that showed regional specificity (**Supp Table 14**). In the cortex and hippocampus, core ion channel proteins included voltage gated calcium channel subunits Cacna1e, Cacna2d3, and NMDA receptor subunit Grin1. Cerebellar Camk2a neuronal signatures were characterized by higher levels of voltage-gated calcium channel subunits Cacna1a, Cacna2d2, Cacng2, chloride channel Clic1, and several potassium channels including Kcnc1, Kcnc3, Kcnd2, Kcnip1. In contrast, pons/medulla Camk2a neurons were characterized by higher expression of potassium channels Hcn2, Kcna2 and Kcnj10. Similar to the AAV-CIBOP results, we identified several (*n* = 52) druggable target proteins that exhibited region-specific patterns (**Supp Table 13**). Drug targets with high levels in cortical Camk2a neurons included ion channel Gria3, a regulator of neurogenesis (Ntrk2), and lipid metabolic protein Plcb1. In contrast, drug targets in the cerebellum included mRNA transport and mRNA processing proteins (e.g., Snrnp200) and the phospholipase Plcb4.

Region specific proteins were further searched against curated lists of gene-disease associations (DisGeNet) to define regional gene-disease associations (**Supp Table 16**). Cortex and hippocampal Camk2a neuron proteins (e.g., Lingo1, Homer1, Gria1) were associated with mental health disorders (schizophrenia, bipolar disease, depression, autism), neurodevelopmental disorders, essential tremor, epileptic encephalopathy, and primary epileptic disorders (e.g., West syndrome). Pons/medulla enriched proteins were associated with dystonia and Parkinson’s disease while proteins enriched in the cerebellum were linked to primary lateral sclerosis, autism, epilepsy, mood disorders, and psychotic disorders.

Since the above analyses were targeted towards proteins with region-specific enrichment patterns, we also analyzed our data in an unbiased manner (K-means clustering). As a result, we identified five clusters of proteins that showed regional patterns of protein expression (**Supp Fig 6a, Supp Table 17**) consistent with region-specific analyses described above. GSEA of the clusters revealed that cortical and hippocampal Camk2a neurons expressed higher levels of proteins involved in GABA signaling and glutamate signaling pathways, neuron development, and synaptic function (Cluster 2 & 4, **Supp Fig 6b, Supp Table 18**) while the cerebellar Camk2a neuronal proteome showed higher levels of proteins involved in hydrolase activity and translation initiation factor 3 (Cluster 5, **Supp Fig 6b, Supp Table 18**). The pons/medulla Camk2a proteome showed a unique signature of pigment granule as well tau-kinase activity related proteins. GSEA also indicated regional metabolic differences in Camk2a neurons, with over-representation of glycolytic, amino acid, and fatty acid metabolic proteins in the pons/medulla (Cluster 1 & 3, **Supp Fig 6b, Supp Table 18**). Intriguingly, we did not identify a cluster distinct to the striatum/thalamus.

In sum, Tg-CIBOP allowed us to comprehensively characterize the proteome of a neuronal sub-type (Camk2a neurons) in the adult mouse brain and resolve regional differences in proteomic composition of Camk2a neurons, which was not easily attainable with the AAV approach or from mouse post-natal or embryonic neuronal culture systems. Importantly, we uncovered potential links between regional proteomic characteristics of Camk2a neurons and disease vulnerability. For example, Camk2a neuronal proteins within the pons/medulla were linked to polyneuropathies and inborn errors in metabolism consistent with the high metabolic protein levels in this region. Cerebellar Camk2a neuronal proteins enriched in mRNA processing and splicing proteins were linked to ataxias, suggesting that splicing dysregulation in cerebellar Purkinje neurons may underlie pathogenesis of cerebellar ataxias. Cortical and hippocampal Camk2a neuronal proteins, enriched in glutamatergic transmission proteins and long-term potentiation, were linked to epilepsy and mood and psychotic disorders emphasizing the relevant of dysfunction in glutamatergic transmission in neuropsychiatric disorders.

### Regional Camk2a neuron-derived phospho-protein signaling and cytokine signatures in adult mouse brain

MS based quantitative proteomics provide a comprehensive and unbiased molecular snapshot of the cell; although, these approaches often fail to detect small and less-abundant proteins such as cytokines and signaling proteins without prior enrichment. Thus, we extended our studies to measure specific biotinylated cytokines and phospho-proteins involved in cellular signaling cascades (e.g., MAPK and Akt/mTOR) using an antibody-based approach. We adapted the multiplexed Luminex sandwich ELISA approach (**Fig 4a**) to directly measure biotinylated phospho-proteins from the MAPK and Akt/mTOR pathways and an array of 32 inflammatory cytokines in brain tissues from both *Rosa26^TurboID/wt^* AAV and transgenic cohorts (**Fig 4b**). First, we captured the proteins of interest onto beads using target-specific antibodies, then a second biotinylated detection antibody followed by a streptavidin fluorophore and fluorescence quantitation (**Fig 4a**, standard assay). In parallel, we adapted this assay to excludes the second biotinylated detection antibody to directly detect neuronal biotinylated proteins of interest via streptavidin fluorophore (**Fig 4a**, adapted assay). This provided an estimate of total and neuron-derived levels of the target protein without the need for enrichment of biotinylated proteins.

**Figure 4.**
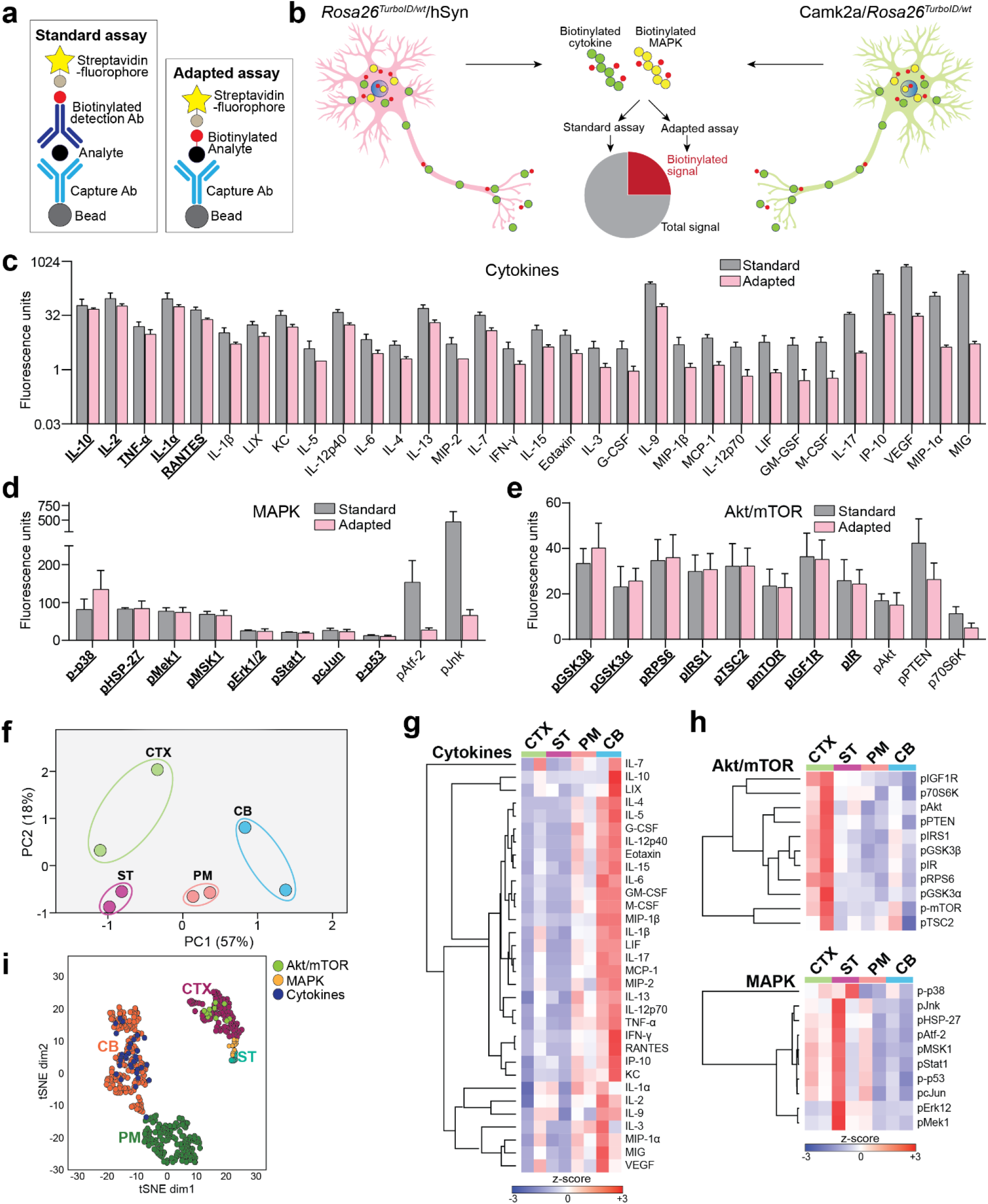
Neuron-specific regional differences in cellular signaling and cytokine levels. **a** Cartoon showing the standard Luminex method to detect total levels of specific proteins and the adapted method to directly measure biotinylated proteins of interest. **b** Cartoon representation of application of standard and adapted Luminex approaches to measure levels of cytokines as well as MAPK and Akt/mTOR pathway phospho-proteins, using whole brain lysates from *Rosa26^TurboID/wt^* AAV (left) and transgenic (right) cohorts. As shown, the readout from the adapted assay represents a subset of the total signal measured in the standard assay. **c, d, e** Fluorescence values (after background subtraction) for cytokines (d), MAPK phospho-proteins (d) and Akt/mTOR phospho-proteins (e) measured using standard (gray) and adapted (pink) assays, from *Rosa26^TurboID/wt^*/AAV9-hSyn-Cre mice (*n* = 3). Predominantly neuron-derived cytokines or phospho-proteins (indicated by bold and underline) have similar fluorescence values in both standard and adapted assays, while predominantly non-neuronal cytokines or phospho-proteins have a markedly lower adapted assay readout as compared to the standard assay. Y-axis shows fluorescence values in a logarithmic scale and error bars represent standard error of mean. **f** Principal component analysis (PCA) of adapted Luminex assays performed on whole brain lysates from the *Rosa26^TurboID/wt^* transgenic cohort, showing regional differences (CTX, ST, PM, and CB) based on measured cytokines and phospho-proteins from the MAPK and Akt/mTOR pathways. Hippocampal lysate was not available for Luminex studies. For this analysis, fluorescence values from adapted assays from labeled (*Rosa26^TurboID/wt^*/Camk2a-Cre^Ert2^) and non-labeled controls (Camk2a-Cre^Ert2^) brain lysates were first normalized to TurboID abundance from MS studies (Figure 3) after which noise (signal from adapted assays from control mice) was then subtracted. **g, h** Heat map showing results from HCA of normalized data from adapted Luminex assays. Analyses of cytokines (g) and phospho-proteins from MAPK and Akt/mTOR pathways (h) are shown, revealing region specific signatures. Overall, neuron-derived cytokines had higher levels in the cerebellum and relatively lower levels in the cortex while phospho-proteins, particularly from the Akt/mTOR pathway had higher levels in the cortex and lower levels in the cerebellum. Related raw and normalized Luminex data from all animals from AAV and transgenic cohorts are included in Supplemental Tables 19-21. **i** An integrated tSNE of core regional proteomic signatures and Luminex data showed clustering of Akt/mTOR signaling with cortex-specific proteins while elevated cytokines clustered with the cerebellar proteomic signature and MAPK clustered with striatal/thalamic proteomic signature.

In the AAV cohort, the standard Luminex assay showed that AAV9-hSyn-Cre increased levels of signaling phospho-proteins (pAkt, pPTEN) and inflammatory cytokines (IL-2, IL-13, IL-17, IP-10, KC and MIG) indicating an inflammatory response related to AAV9 injection (**Supp Table 19**). Given this effect of AAV9, we subsequently compared *Rosa26^TurboID/wt^*/hSyn mice to WT/hSyn mice using the standard assay and found significant elevation of 10 phospho-proteins (e.g., pMSK1, pStat1, pMek1) and 9 cytokines (e.g., VEGF, MIG, IL7) in labeled mice. This increased phospho-protein and cytokine response in *Rosa26^TurboID/wt^*/hSyn mice could be due to either an inflammatory response or due to additive increased signal of biotinylated target proteins in the assay. Despite these differences between groups using the standard assay, the adapted Luminex approach detected 9-fold higher signal in the *Rosa26^TurboID/wt^*/hSyn brains compared to minimal signal in the control brains, confirming that the elevated signal was indeed due to biotinylation by TurboID. By comparing the standard assay signal (total analyte abundance) with the adapted assay signal (biotinylated neuron-derived signal) in the *Rosa26^TurboID/wt^*/hSyn brain lysates, we found that majority of cytokines, with the exceptions of IL-10, IL-2, TNF-α, IL-1α, and RANTES (**Fig 4c**, underlined proteins), were not neuronally derived. Remarkably, we quantified several MAPK phosphoproteins (**Fig 4d**, underlined proteins, e.g., p-p38, pMek1 and pErk1/2) and Akt/mTOR phosphoproteins (**Fig 4e**, underlined proteins, e.g., pGSK-α and −β, p-mTOR) as being predominantly neuronally derived.

We also applied the adapted Luminex approach to investigate regional differences in neuron-derived signaling proteins and cytokines in *Rosa26^TurboID/wt^*/Camk2a mice. After accounting for background, adapted Luminex assay signals were negligible in the control mice (**Supp Table 20**). Highest signal intensities in the adapted assay across all regions were observed for 25 of 32 cytokines (Top 5: IL-2, IL-1α, IL-9, IP-10, and VEGF) and phospho-proteins from both Akt/mTOR (11 of 11 analytes) and MAPK (10 of 10) signaling pathways. Similar to the results from our AAV cohort, phosphoproteins from signaling pathways were more robustly biotinylated as compared to cytokines (**Supp Table 21**). A combined PCA of fluorescence intensities from adapted Luminex assays for MAPK, Akt/mTOR, and cytokine panels revealed four distinct clusters associated with brain regions of *Rosa26^TurboID/wt^*/Camk2a mice (**Fig 4f**). Specifically, we identified unique patterns of neuron-derived cytokine levels that distinguished the cerebellum from the cortex and pons/medulla after adjusting for TurboID levels (**Fig 4g**). Intriguingly, cortical Camk2a neuron-derived signature included low levels of cytokines but high levels of phospho-proteins from the Akt/mTOR pathway (**Fig 4h**). In contrast, cerebellar Camk2a neuronal signature displayed higher level of cytokines and low levels of Akt/mTOR and MAPK phospho-proteins, while MAPK phospho-proteins appear to be enriched in the striatum/thalamus. (**Fig 4g-h**). An integrated t-distributed stochastic neighbor embedding (tSNE) analysis of core regional proteomic signatures derived from LFQ-MS studies and from Luminex data confirmed clustering of Akt/mTOR signaling with cortex-specific proteins while elevated cytokines clustered with the cerebellar proteomic signature (**Fig 4i**).

Collectively, these targeted immunoassays of biotinylated proteins provide a complimentary approach to MS for detecting proteomic signatures that resolve signaling mechanisms underlying cell-type specific phenotypes in brain.

## DISCUSSION

We report a novel cell type-specific *in vivo* proteomic labeling approach in which the biotin ligase, TurboID, is expressed in a desired cell type using a Cre-lox genetic strategy. We have generated and validated the novel *Rosa26^TurboID^* mouse line that can now be bred with a plethora of well-validated existing non-inducible and inducible Cre mouse lines to achieve tissue as well as cell type-specific proteomic labeling. The broad cellular proteomic labeling allows us to capture a snapshot of the cellular proteome of a specific cell type while retaining the native state of the cell since the labeled proteins can be purified directly from brain lysates without cell type isolation. We term this approach cell type-specific *in vivo* biotinylation of proteins (CIBOP). To demonstrate the utility of CIBOP, we successfully performed proof-of-principle experiments to label the neuronal proteome using AAV and genetic approaches, and then used unbiased MS and targeted immunoassays of neuron-derived biotinylated proteins from whole brain samples. LFQ-MS allowed us to identify over 2,000 neuronal proteins and novel regional differences within Camk2a neurons. A major advantage of the TurboID approach is its depth of proteomic coverage, even with ‘single-shot” label-free MS approaches. We anticipate 3-4-fold greater proteomic coverage in future studies with tandem multiplexed approaches coupled with off-line fractionation^25^. While we did not specifically perform neuron-specific phospho-proteomic profiling of neurons in our studies, we were able to quantity over 140 phosphoproteins without enrichment. This depth of phosphoprotein coverage could be increased several-fold with phosphopeptide enrichment approaches^26^ which could reveal novel cell type-specific proteomic insights at the level of post-translational modifications that cannot otherwise be captured by other methods. Our successful application of CIBOP to investigate proteomic profiles of neurons in their native state, demonstrates the validity of this approach for broad applications in neuroscience and outside the nervous system, by leveraging the plethora of conditional Cre/lox models that are currently available for cell type and tissue-type manipulation in mouse models.

Since CIBOP involves biotinylation of lysine residues, it is important to consider potential implications of excessive biotinylation on protein and cellular function. Biotin is a water soluble, readily cell permeable, and a brain penetrant vitamin (vitamin H or B7) that is required as a co-factor for several enzymes involved in glucose, amino acid, and fatty acid metabolism^27^. While biotin deficiency can cause growth retardation, skin and neurological diseases, high doses of systemically administered biotin are non-toxic^21,28^. Since TurboID is highly efficient, biotin supplementation is necessary to prevent a relative biotin deficiency caused by excessive protein biotinylation which may shunt biotin away from biotin-dependent cellular processes^18,29^. Appropriately, some concerns regarding toxicity of excessive biotinylation have been raised in the literature^18^. In the present study, we did not observe any adverse effects of neuron-specific biotinylation in *Rosa26^TurboID^* mice, supported by electrophysiological studies. Whether shortening the duration and/or degree of biotinylation can still achieve sufficient proteomic labeling needs to be determined. As CIBOP is extended to other cell types, investigators will need to optimize the dose and duration of biotin supplementation to individual applications.

The biotinylated proteomes from AAV and transgenic CIBOP approaches were highly enriched with proteins involved in neurotransmission and axon guidance, consistent with neuron-specific labeling and widespread neuronal cell body, axonal, and dendritic biotinylation with no glial labeling or activation. We captured synaptic proteins, ion channels, and membrane receptors including druggable targets. AKT1, a signaling protein which is a key member of the IGF1/PI3K/AKT1/mTOR axis that is relevant to synaptic functioning and memory, is a druggable target that was identified in the AAV and transgenic proteomes, highlighting consistency across both CIBOP datasets and emphasizing important roles for this pathway in neuronal physiology. At the sub-cellular level, cytosolic and membrane associated proteins such as ion channels, as well as organelles such as mitochondria, ER and Golgi apparatus, and vesicles, were abundant in the enriched neuronal proteome. We attribute the broad proteomic labeling to a combination of the biotinylation radius (≈10nm) of biotin ligases^19^, the promiscuous nature of protein biotinylation by TurboID, and the use of a nuclear export sequence (NES) in the transgene of *Rosa26^TurboID^* mice. The labeling of secreted and vesicular proteins also suggests that simultaneous profiling of secreted or released proteins in tissues and biofluids may be feasible. Indeed, secretome profiling using *in vivo* TurboID via AAV delivery was successful in a non-brain context ^30^. Our ongoing studies with *Rosa26^TurboID^* mice, in brain and non-brain disease contexts, will determine whether cellular origin of secreted proteins can indeed be measured in biofluids as biomarkers of underlying cellular mechanisms of disease.

The Tg-CIBOP strategy also successfully labeled neurons in the adult brain and, regardless of brain region, the proteome showed enrichment of neuronal synaptic proteins such as Dlg4 (PSD-95), Gap43, Rph3a, and Vamp2. With this approach, we had the opportunity to characterize regional proteomic differences within the same neuronal sub-type and found core protein signatures unique to the cortex and hippocampus, striatum/thalamus, pons/medulla, and cerebellum while overcoming limitations associated with AAV-CIBOP. Glutamatergic synaptic transmission, calcium signaling, and synaptic plasticity encompassed the proteomic signature of cortical and hippocampal Camk2a neurons. Alternatively, cerebellar proteomic signatures point to mRNA processing, splicing, and calcium transporting ATPase activity. Unique proteomic signatures of pontine and medullary neurons with increased expression of metabolic proteins, inflammatory proteins, and pigmented granules agrees with the expected enrichment of axonal tracks and islands of neuronal cell bodies in the brain stem, particularly pigmented neurons^31^. Collectively, these data emphasize the diverse functions of proteins within Camk2a neurons and how each brain region may be functionally distinct from the other. Furthermore, these functionally distinct characteristics reflect neuronal-specific mechanisms within brain regions that may be vulnerable to neurological diseases.

Indeed, several protein/gene-disease associations were enriched among core regional protein lists. In the cortex and hippocampus Camk2a neurons, we identified signature proteins linked to epilepsy (e.g., Dlg4) and schizophrenia (e.g., Grin2a). Several studies have shown that the *GRIN2A* gene, which encodes a glutamate (N-methyl-D-aspartic acid [NMDA]) receptor subunit protein, contains a polymorphism in the promoter that is predisposes individuals to schizophrenia^32–35^. Grin2a is also known to interact directly and indirectly with Dlg4 (PSD-95), which regulates glutamate NMDA and α-amino-3-hydroxy-5-methyl-4-isoxazoleproprionic acid (AMPA) receptor trafficking at the synapse^36^ with significant roles in Alzheimer’s disease (AD) biology^37^. Interestingly, regional differences in phospholipase C isoenzyme (Plcb1, Plcb4), druggable targets identified in the Camk2a proteome, expression in the brain are known to underlie several brain disorders including epilepsy, schizophrenia, and AD^38^. The neurotrophic tyrosine kinase receptor type 2 (NTRK2), a protein significantly enriched in our cortical and hippocampal Camk2a proteome associated with mood disorders, is a specific receptor for neurotrophin brain-derived neurotrophic factor (BDNF) which is implicated in the pathophysiology of bipolar disorder^39^. In the cerebellum, core signature proteins were expectedly associated with ataxias, which are defined by Purkinje cell and cerebellar circuitry dysfunction^40^. One such protein we identified is Cacna1a, a α1A-subunit of voltage-gated P/Q-type calcium channels (Cav2.1), which contains a mutation that causes spinocerebellar ataxia type 6 (SCA6) and episodic ataxia type 2^41,42^. Lastly, proteins related to inborn errors of metabolism (e.g., Acadsb)^43^ and inherited polyneuropathies (e.g., Nefl) were highly abundant in the pons/medulla. Interestingly, dominant mutations in *NEFL* cause clinically distinct forms of Charcot-Marie-Tooth disease (CMT)^44–47^, ranging from a severe neuropathy with an infantile onset, to a more moderate neuropathy with onset typically between 10 and 20 years. In summary, these regional protein/gene-disease associations signify the ability of CIBOP to identify mechanisms underpinning selective neuronal and regional vulnerability as well as its potential application to specific disease models to resolve the molecular basis of this vulnerability temporally in various neurological diseases^48^.

In our CIBOP studies, we found that several MAPK (e.g., pERK1/2 and pMEK1/2) and Akt (e.g., pGSK3β and pGSK3α phosphoproteins that are highly abundant in neurons while majority of cytokines in the brain, with the exceptions of IL-10, IL-2, TNF-α IL-1α and RANTES were mostly of non-neuronal origin. Previous studies have shown that neurons may produce cytokines such as IL-2 and TNF-α under homeostatic conditions^49–52^. We also resolved novel differences in neuronal MAPK and Akt signaling activation across brain regions, with cortical Camk2a neurons exhibiting the highest level of baseline activation Akt/mTOR signaling and striatal/thalamic neurons exhibiting highest level of MAPK. From a disease standpoint, recent studies have shown that increased MAPK protein expression and MAPK pathway activation (e.g., ERK signaling) are highly characteristic and potentially causal, in neurodegenerative diseases such as AD^53,54^. MAPK signaling is also critical for synaptic mechanisms such as long-term potentiation and response to stress or injury^53^. Akt signaling activates the mTOR pathway that regulates mRNA translation, metabolism, and protein turnover, and is highly relevant to the pathogenesis of lysosomal storage disorders, genetic neurodevelopmental disorders, and neurodegenerative disorders^55^. The ability of CIBOP to label and quantify neuron-specific MAPK and Akt signals from whole brain can be highly relevant to understanding the spatio-temporal dynamics of these signaling pathways in disease pathogenesis. While our observed regional differences in neuron-derived cytokines are exploratory, the inverse correlation between MAPK and Akt signaling (high in the cortex but low in cerebellum) and cytokine levels (low in the cortex but high in the cerebellum), warrant further investigation^56^.

## CONCLUSION

We have generated and validated a novel *Rosa26^TurboID^* mouse line that enables CIBOP, a powerful experimental approach to investigate molecular changes occurring at the proteomic level. Data obtained using this *in vivo* system showcases breadth of proteomic labeling in neurons, regional Camk2a neuronal proteomic signatures, and neuron-specific phospho-protein signaling (MAPK and Akt) and cytokine signatures that are associated with various neurological disorders. This novel mouse model and our validated experimental workflow provide a framework to resolve cellular mechanisms underlying physiological or pathological conditions of complex tissues.

## METHODS

### Construct generation and cell culture studies

The V5-TurboID-NES_pCDNA3, a gift from Alice Ting^18^ (Addgene plasmid # 107169) was used to generate AsiS1-Kozak-V5-TurboID-NES-stop-Mlu1 construct. This was then cloned into the pR26 CAG AsiSI/MluI targeting vector, a gift from Ralf Kuehn^57^ (Addgene plasmid # 74286), to generate the a Rosa26 (chromosome 6) targeting vector containing the CAG promoter, a floxed STOP site (loxp-STOP-loxp), and V5-TurboID with a nuclear export signal (TurboID-NES). This Rosa26^TurboID^ targeting vector was verified *in-vitro* for Cre-mediated TurboID expression and biotinylation in HEK293 cells (**Supp Fig 1a-b**).

Human embryonic kidney 293 (HEK293) cells were maintained in Dulbecco’s modified Eagle’s medium (DMEM) supplemented with 10% fetal bovine serum (FBS) and 1% penicillin/streptomycin at 37°C in a 5% CO_2_ atmosphere. For transient transfection, cells grown to 70-80% confluency in 6-well plates were transfected (Lipofectamine 3000, Thermo, L3000001) with 2.5 µg/well of Rosa26^TurboID^ targeting vector and Cre plasmid (CMV-Cre, a gift from Dr. Xinping Huang, Emory University), Rosa26^TurboID^ targeting vector alone, or Cre plasmid alone according to manufacturer’s protocol. Untransfected cells also served as negative controls. Twenty-four hours post-transfection, cells were treated with 200 μM biotin for another 24 hrs. Subsequently, cells were rinsed with cold 1X phosphate buffered saline (PBS) and harvested in urea lysis buffer (8 M urea, 10 mM Tris, 100 mM NaH_2_PO_4_, pH 8.5) containing 1X HALT protease inhibitor cocktail without EDTA (Thermo, 87786). The cells were sonicated for 3 rounds consisting of 5 sec of active sonication at 25% amplitude with 10 sec incubation periods on ice between sonication. Lysed cells were then centrifuged for 5 min at 15,000 × g and the supernatants were transferred to a new tube. Protein concentration was determined by bicinchoninic acid (BCA) assay (Thermo, 23225). To confirm protein biotinylation, 10 μg of cell lysates were resolved on a 4-12% Bris-Tris gel, transferred onto a nitrocellulose membrane, and probed with streptavidin-Alexa680 (Thermo, S32358) diluted 1:10K in Start Block (Thermo, 37543) for one hour at room temperature. Subsequently, the membrane was washed in 1X tris buffered saline containing 0.1% Tween20 (TBS-T) and incubated over-night with mouse anti-V5 (1:250; Thermo, R960-25) or anti-Gapdh (1:3000; Abcam, ab8245) diluted in Start Block. After washes in TBS-T, membranes were incubated with goat anti-mouse 700 (1:10K) and imaged using Odyssey Infrared Imaging System (LI-COR Biosciences).

For immunocytochemistry (ICC), cells grown to 50% confluency on coverslips were transfected and treated with biotin as described above. Subsequently, cells were washed 3X with warm sterile PBS for 5 min each and fixed with 1X ICC fixation buffer (Invitrogen, 00-8222-49) for 30 min. After PBS washes, cells were permeabilized (Invitrogen, 00-8333-56) for 20 min and blocked in 10% normal horse serum (NHS) in PBS for 45 min at room temperature. The cells were then incubated with mouse anti-V5 diluted in 2% NHS in PBS overnight. After thorough washes with PBS, cells were incubated with anti-mouse Rhodamine Red (1:500) and streptavidin Alexa-flour 488 (1:500, Thermo, S11223) diluted in 2% NHS in PBS for 1 hour. Coverslips were mounted onto slides using mounting media containing DAPI (Sigma-Aldrich, F6057) and images were captured using an Olympus fluorescence microscope (Olympus BX51) and camera (Olympus DP70) and processed using Image J software (FIJI Version 1.51).

### Generation of TurboID-floxed mouse

The Rosa26^TurboID^ targeting vector was electroporated into JM8A3 C57BL/6N embryonic stem (ES) cells. After confirming homologous recombination in ES clones, they were microinjected into albino goGermline^TM^ blastocysts (Ozgene) and implanted into pseudo pregnant CD-1 female mice^21^ (Emory Mouse Transgenic and Gene Targeting Core). The resulting chimeric mice were crossed to wild-type (WT) C57BL6/J^58^ mice to yield F1 *Rosa26^TurboID^* heterozygous mice (*Rosa26^TurboID/wt^*) and littermate controls. Genotyping was performed on DNA extracted from a tail biopsy from mice using the following primers: (TurboID_fwd) 5’ ATCCCGCTGCTGAACGCTAAAC 3’, (TurboID_rev) 5’ ACCATTTCCTCCCTCTGCTTCC 3’, (ROSA26_fwd) 5’ CTCTTCCCTCGTGATCTGCAACTCC 3’, (ROSA_rev) 5’ CATGTCTTTAATCTACCTCGATGG 3’. Mice were designated as heterozygous (*Rosa26^TurboID/wt^*) by the presence of a band approximately 181bp corresponding to the *TurboID* transgene and a 299bp band produced by the endogenous *ROSA26* allele, while WT contained only the 299bp endogenous *ROSA26* band (**Supp Fig 1c**).

### Animal studies

All mice used in the present study were housed in the Department of Animal Resources at Emory University under a 12-hour light/12-hour dark cycle with ad libitum access to food and water. All procedures were approved by the Institutional Animal Care and Use Committee of Emory University and were in strict accordance with the National Institute of Health’s “Guide for the Care and Use of Laboratory Animals”.

AAV approach for Cre recombinase delivery to neurons in *Rosa26^TurboID/wt^* mice: Two-month-old mice were anesthetized with isoflurane, given sustained-release buprenorphine subcutaneously (0.5 mg/kg), and immobilized on a stereotaxic apparatus. The mice were maintained on 1% - 2.5% isoflurane and monitored closely for breathing abnormalities throughout the surgery. Bilateral intrahippocampal injections (coordinates from bregma: −2.1 mm posterior, ±2.0 mm lateral, and ±1.8 mm ventral) were performed over a 5 min period with 1 µL of AAV9-hSyn-Cre (Titer ≥ 1×10¹³ vg/mL, Addgene, Cat. No. 105553-AAV9) or un-injected: un-injected WT mice (*n* = 3, male), wild-type mice injected with AAV9-hSyn-Cre (*n* = 3, male), and *Rosa26^TurboID/wt^* mice injected with AAV9-hSyn-Cre (*n* = 3, male) (**Fig 1b**). The incision was closed with tissue adhesive (Fisher, NC0304169), isoflurane was discontinued, and the animal was revived in a new, clean cage atop a heating pad. The mice were monitored every 15 min for 1 hour after surgery and routinely for the 3-day post-surgery survival period under normal vivarium conditions. After 4 weeks, mice were given water supplemented with biotin^21^ (37.5 mg/L)^21^ for two weeks until euthanasia at 3 months of age.

Transgenic cohort: *Rosa26^TurboID/wt^* mice were crossed with Camk2a-Cre^Ert2^ mice (Jackson Labs, Stock No. 012362) to obtain Camk2a-Cre^Ert2^ (“Camk2a”, *n* = 2; male) and *Rosa26^TurboID/ wt^*/Camk2a-Cre^Ert2^ (“*Rosa26^TurboID/wt^*/Camk2a”, *n* = 2; 1 male and 1 female) littermate mice. All mice were given tamoxifen (75mg/kg) intraperitoneally for five days at 6 weeks of age. After 3 weeks, mice were given water supplemented with biotin (37.5mg/L)^21^ for two weeks until euthanasia at 3 months of age. Consistent with previous publications^24^, we have previously validated the Camk2a-cre^Ert2^ line for neuron-specific labeling and non-leaky Cre activity.

After biotin supplementation, mice were anesthetized with ketamine (ketamine 87.5 mg/kg, xylazine 12.5 mg/kg) followed by transcardial perfusion with 30mL of ice-cold PBS. The brain was immediately removed and hemi-sected along the mid-sagittal line. The left hemisphere was fixed in 4% paraformaldehyde (PFA) for 24 hours and then transferred to 30% sucrose after through washes in PBS. The right hemisphere was either immediately snap frozen on dry ice for the AAV cohort or dissected for the following brain regions for the transgenic cohort and then snap frozen: cortex (CTX), hippocampus (HIP), cerebellum (CB), pons/medulla (PM), and striatum/thalamus (ST) (**Fig 2a**).

### Tissue homogenization and immunoblotting

Frozen brain tissue pieces (AAV cohort: forebrain without CB and PM; transgenic cohort: dissected brain regions) were added to 1.5 mL Rino tubes (Next Advance) containing stainless-steel beads (0.9-2 mm in diameter) and five volumes of the tissue weight in urea lysis buffer (8 M urea, 10 mM Tris, 100 mM NaH_2_PO_4_, pH 8.5) containing 1X HALT protease inhibitor cocktail without EDTA. Tissues were homogenized in a Bullet Blender (Next Advance) twice for 5 min cycles at 4 °C. The homogenates were transferred to a new Eppendorf LoBind tube followed by 3 rounds of sonication consisting of 5 sec of active sonication at 25% amplitude with 10 sec incubation periods on ice between sonication. Homogenates were then centrifuged for 5 min at 15,000 × g and the supernatants were transferred to a new tube. Protein concentration was determined by BCA assay.

To confirm protein biotinylation, 20 μg of brain lysates were resolved on a 4-12% Bris-Tris gel, transferred onto a nitrocellulose membrane, and probed with streptavidin-Alexa 680 diluted in Start Block for 1 hour at room temperature. Subsequently, the membrane was incubated over-night with mouse anti-V5 or anti-Gapdh diluted in Start Block. After washes in TBS-T, membranes were incubated with goat anti-mouse 700 (1:10K) and imaged using Odyssey Infrared Imaging System (LI-COR Biosciences). Biotinylation was quantified using Odyssey Studio Lite. For densitometry, a box around the entire length of streptavidin signal was drawn for every lane and normalized to Gapdh signal.

### Biotinylated protein enrichment

After confirmation of biotinylation of proteins and V5 expression by Western blot, biotin-tagged proteins were enriched using a previously published protocol^19^ with slight modifications. Briefly, for each sample, 83 μL of streptavidin magnetic beads (Thermo, 88817) in a 1.5 mL Eppendorf LoBind tube were washed twice with 1 mL RIPA lysis buffer (50 mM Tris, 150 mM NaCl, 0.1% SDS, 0.5% sodium deoxycholate, 1% Triton X-100) on rotation for 2 min. The beads were incubated at 4 °C for at least 1 hour with rotation with 500 µg - 1 mg protein from each sample with an addition 500μL RIPA lysis buffer. The beads were centrifuged briefly, placed on a magnetic rack, and the supernatant was collected into a new 1.5 mL Eppendorf LoBind tube and frozen at −20 °C. The beads were washed in the following series of buffers on rotation at room temperature: twice with 1 mL RIPA lysis buffer for 8 min, once with 1 mL 1 M KCl for 8 min, once with 1 mL 0.1 M sodium carbonate (Na_2_CO_3_) for ∼10 sec, once with 1mL 2M urea in 10 mM Tris-HCl (pH 8.0) for ∼10 sec, and twice with 1 mL RIPA lysis buffer for 8 min. The beads, in the final RIPA lysis buffer wash, were transferred to a new tube and washed twice with 1 mL of PBS for 2 min on rotation. After the final wash, beads were resuspended in 83 μL of PBS. To verify protein enrichment, 10% of the bead volume was transferred to a new tube and were boiled in 30 μL of 2X protein loading buffer (Biorad, 1610737) supplemented with 2 mM biotin and 20 mM dithiothreitol (DTT) at 95 °C for 10 min to elute the biotinylated proteins. Subsequently, 10 μL of eluate was run on a gel and probed with Streptavidin680 while 20 μL of the eluate was run on a separate gel and Silver stained (Thermo, 24612). All samples in the *Rosa26^TurboID/wt^* AAV cohort (*n* = 3/condition, *n*_total_ = 9) were enriched as described above. For the transgenic cohort (*n* = 2/brain region/genotype), we enriched from 500 µg of protein and prepared technical replicates for enrichment from the *Rosa26^TurboID/wt^*/Camk2a brain regions, which resulted in *n* = 4/region, while the control Camk2a brain regions did not contain technical replicates (**Fig 2a**).

### Mass spectrometry

To prepare enriched samples for mass spectrometry analysis, the remaining 90% of streptavidin beads were washed three times with PBS and then resuspended in 50 mM ammonium bicarbonate (NH_4_HCO_3_). Bound proteins were then reduced with 1 mM dithiothreitol (DTT) at room temperature for 30 min and alkylated with 5 mM iodoacetamide (IAA) in the dark for 30 min with rotation. Proteins were digested overnight with 0.5 µg of lysyl endopeptidase (Wako) at RT on shaker followed by further overnight digestion with 1 µg trypsin (Thermo, 90058) at RT on shaker. The resulting peptide solutions were acidified to a final concentration of 1% formic acid (FA) and 0.1% triflouroacetic acid (TFA), desalted with a HLB columns (Waters, 186003908) and dried down in a vacuum centrifuge (SpeedVac Vacuum Concentrator). Similarly, 50 µg of protein from pooled total brain lysates from the AAV cohort were digested and desalted.

Dried peptides were resuspended in 15 μL of loading buffer (0.1% FA and 0.03% TFA in water), and 7-8 μL was loaded onto a self-packed 25 cm (100 μm internal diameter packed with 1.7 μm Water’s CSH beads) using an Easy-nLC 1200 or Dionex 3000 RSLCnano liquid chromatography system. The liquid chromatography gradient started at 1% buffer B (80% acetonitrile with 0.1% FA) and ramps to 5% in 10 seconds. This was followed by a 55 min linear gradient to 35% B and finally a 4 minute 50 second 99% B flush. For the AAV cohort (IP and total brain samples), an Orbitrap Lumos Tribrid mass spectrometer with a high-field asymmetric waveform ion mobility spectrometry (FAIMS Pro) interface was used to acquire all mass spectra at a compensation voltage of -45V. The spectrometer was operated in data dependent mode in top speed mode with a cycle time of 3 seconds. Survey scans were collected in the Orbitrap with a 60,000 resolution, 400 to 1600 m/z range, 400,000 automatic gain control (AGC), 50 ms max injection time and rf lens at 30%. Higher energy collision dissociation (HCD) tandem mass spectra were collected in the ion trap with a collision energy of 35%, an isolation width of 1.6 m/z, AGC target of 10000, and a max injection time of 35 ms. Dynamic exclusion was set to 30 seconds with a 10 ppm mass tolerance window.

For the transgenic brain region IP samples, an Orbitrap Eclipse Tribrid mass spectrometer with a high-field asymmetric waveform ion mobility spectrometry (FAIMS Pro) interface was used to acquire all mass spectra at a compensation voltage of -45V. The spectrometer was operated in data dependent mode in top speed mode with a cycle time of 3 seconds. Survey scans were collected in the Orbitrap with a 120,000 resolution, 400 to 1600 m/z range, 400,000 automatic gain control (AGC), 50 ms max injection time and rf lens at 30%. Higher energy collision dissociation (HCD) tandem mass spectra were collected in the ion trap with a collision energy of 35%, an isolation width of 0.7 m/z, AGC target of 10000, and a max injection time of 35 ms. Dynamic exclusion was set to 30 seconds with a 10 ppm mass tolerance window. The mass spectrometry raw data have been deposited to the ProteomeXchange Consortium via the PRIDE partner repository with the dataset identifier: px-submission #519326.

### Protein identification and quantification

MS raw files were searched using the search engine Andromeda, integrated into MaxQuant, against 2020 mouse Uniprot database (91,441 target sequences including peptide sequences for V5 and TurboID). Methionine oxidation (+15.9949 Da) and protein N-terminal acetylation (+42.0106 Da) were variable modifications (up to 5 allowed per peptide); cysteine was assigned as a fixed carbamidomethyl modification (+57.0215 Da). Only fully tryptic peptides were considered with up to 2 missed cleavages in the database search. A precursor mass tolerance of ±20 ppm was applied prior to mass accuracy calibration and ±4.5 ppm after internal MaxQuant calibration. Other search settings included a maximum peptide mass of 4,600 Da, a minimum peptide length of 6 residues, 0.05 Da tolerance for orbitrap and 0.6 Da tolerance for ion trap MS/MS scans. The false discovery rate (FDR) for peptide spectral matches, proteins, and site decoy fraction were all set to 1 percent. Quantification settings were as follows: re-quantify with a second peak finding attempt after protein identification has completed; match MS1 peaks between runs; a 0.7 min retention time match window was used after an alignment function was found with a 20-minute RT search space. Quantitation of proteins was performed using summed peptide intensities given by MaxQuant. The quantitation method only considered razor plus unique peptides for protein level quantitation.

The MaxQuant output data were uploaded onto Perseus (Version 1.6.15) for analyses. The categorical variables were removed, and intensity values were log (base 2) transformed. The data were filtered to contain at least 3 valid values across 9 samples to accommodate 6 missing values in the negative controls for the AAV cohort and missing values were further imputed from normal distribution (width: 0.3, down shift: 1.8) (**Supp Table 1**). The MaxQuant output data from pooled total brain lysates from the AAV cohort were filtered to contain at least 1 valid value across 3 samples and missing samples were imputed from normal distribution (width: 0.3, down shift: 1.8) (**Supp Table 6**). Lastly, the MaxQuant output data from the transgenic cohort brain regions were first filtered to contain at least 9 valid intensity values in the transgenic IP samples and then processed as described above with Perseus. Afterwards, the abundance values in the transgenic IP samples were normalized to TurboID abundance across all brain regions (**Supp Table 10**).

### Differential expression, Gene Ontology analysis, Clustering, and data visualization

We applied differential expression, gene ontology (GO) analysis, as well as principal component analysis (PCA) and hierarchical clustering approaches (average linkage method, one minus Pearson correlation) to analyze proteomic data from both *Rosa26^TurboID/w^*^t^ AAV and transgenic cohorts. Heat maps of normalized data were generated using Morpheus (Broad Institute, Morpheus, https://software.broadinstitute.org/morpheus) and additional graphical representation of data was performed using SPSS (IBM, Version 26.0), R software, and Prism (Graphpad, Version 9).

For the AAV cohort, significantly differentially enriched proteins (unadjusted *p* value ≤ 0.05) were identified by an unpaired t-test comparing a) *Rosa26^TurboID/wt^*/hSyn vs. both control gourps, b) *Rosa26^TurboID/wt^*/hSyn vs. WT un-injected, or c) *Rosa26^TurboID/wt^*/hSyn vs. WT/hSyn (**Supp Table 1**). Differentially enriched proteins are presented as volcano plots.

For the transgenic cohort, prior to any analyses, protein abundances in Camk2a/*Rosa26^TurboID/wt^* brain regions were normalized to TurboID levels to adjust for regional variability. Then, we used an unpaired t-test comparing all brain regions from Camk2a controls vs. all brain regions from *Rosa26^TurboID/wt^*/Camk2a to identify differentially enriched or depleted proteins, shown as a volcano plot (**Supp Fig 5, Supp Table 10**). To identify differentially enriched proteins across brain regions of *Rosa26^TurboID/wt^*/Camk2a mice, we performed an unpaired t-test comparing one brain region vs. all other brain regions (i.e., PM vs. [CTX, HIP, ST, CB]) and displayed proteins that were significantly (*p* ≤ 0.05) ≥4 fold enriched in a heatmap **(Fig 3c, Supp Table 13).**

Gene set enrichment analysis (GSEA) of differentially enriched proteins was performed using AltAnalyze (Version 2.0). GO terms meeting Fisher exact significance of *p* value ≤ 0.05 (i.e., a Z-score greater than 1.96) were considered. Input lists included proteins that were significantly differentially enriched (unadjusted *p* ≤ 0.05, ≥2 fold) from the various differential analyses described above (**Supp Tables 3, 7-9, 12, 14, 15**). The background gene list consisted of unique gene symbols for all proteins identified and quantified in the mouse brain (*n* = 7,736)^5^. For the regional analysis of neuronal proteomic data, we identified proteins that were at least 4-fold enriched in the specific region compared to all other brain regions, representing a core set of regionally enriched proteins. These regional protein lists were searched against curated sets of gene-disease, gene-trait, and gene-phenotype associations (DisGenNet v7.0) and the results relevant to neurological or psychiatric traits were sorted based on gene-disease association scores^59^ (**Supp Table 16**). Networks of known direct (protein-protein) and indirect (functional) interactions within core regional protein signatures were explored using publicly available sources of these interactions (STRING Consortium 2020 Version 11.0)^60^. In order to identify druggable protein targets, we searched our AAV and transgenic proteomic data against a list (*n* = 1,326) of potential drug targets that belong to the following drug target protein classes: enzymes, transporters, receptors. and ion-channels (https://www.proteinatlas.org/humanproteome/tissue/druggable).

### Immunofluorescent staining and microscopy

Fixed brains were cut into 30µm thick sagittal sections using a cryostat. For immunofluorescence (IF) staining, 2-3 brain sections from each mouse were thoroughly washed to remove cryopreservative, blocked in 8% normal horse serum (NHS) diluted in TBS containing 0.1% Triton X-100 for 1 hour, and incubated with primary antibodies diluted in PBS containing 2% NHS overnight (1:500 rabbit anti-Iba1, Abcam, ab178846; 1:200 mouse anti-Map2, BD Pharmagen, 556320; 1:500 mouse anti-Gfap, Thermo, 14-9892-82). Following thorough washes and incubation in the appropriate fluorophore-conjugated secondary antibody (1:500, anti-mouse Rhodamine-red or anti-rabbit Rhodamine-red) and streptavidin Alexa-flour 488 for 1 hour at room temperature, sections were mounted on slides with mounting media containing DAPI (Sigma-Aldrich, F6057) for nuclear staining. Representative images of the same regions across all samples were taken using the Keyence BZ-X810 and all images processing was performed using Image J software (FIJI Version 1.51).

### Quantification of cytokines and signaling phospho-protein levels in brain homogenates

Luminex multiplexed immunoassays were used to quantify phospho-proteins within the MAPK (Millipore 48-660MAG) and PI3K/Akt/mTOR (Millipore 48-612MAG) pathways as well as 32 panel of cytokine/chemokine protein expression (Millipore MCYTMAG-70K-PX32) in brain lysates from the *Rosa26^TurboID/wt^* AAV and transgenic cohorts. Samples were analyzed using standard Luminex assays, consisting of a both capture and biotinylated detection antibody steps, termed “standard assay”, and a customized Luminex protocol in which the biotin detection antibody step was omitted, termed “adapted assay”. Linear range analysis was conducted to identify a range of total protein loaded that corresponded to linear signal readout from the Luminex instrument for each analyte. Based on this, 6 μg, 0.375 μg, and 2 μg of total protein from the AAV cohort was loaded for the cytokine, PI3K/Akt/mTOR, and MAPK assays, respectively. For the brain regions from the transgenic cohort, 2.5 μg, 2 μg, and 0.75 μg of total protein was loaded for the cytokine, PI3K/Akt/mTOR, and MAPK assays, respectively. Assays were read on a MAGPIX instrument (Luminex).

For the *Rosa26^TurboID/wt^* AAV cohort, the background was subtracted from fluorescence intensity values for each analyte from the standard and adapted assays. Data from each assay are shown as bar graphs and are detailed in **Supplemental Table 19**. For the brain regions from the *Rosa26^TurboID/wt^* transgenic cohort, fluorescent intensity values from the adapted Luminex assay from the control mouse samples were subtracted from the intensity values from the adapted assay from *Rosa26^TurboID/wt^*/Camk2a mouse samples. These values were then normalized to TurboID abundance from mass spectrometry intensity for each region in order to account for differences in Camk2a promoter activity and TurboID protein expression. We then performed hierarchical clustering analysis of the normalized data (Morpheus) to identify regional patterns which were visualized as heatmaps (raw and normalized data included in **Supp Tables 20 & 21**).

### Acute hippocampal slice preparation

Acute hippocampal slices were prepared from 3-month-old *Rosa26^TurboID/wt^*/Camk2a mice or littermate control *Rosa26^TurboID/wt^* after receiving tamoxifen and biotin supplementation. Mice were first anesthetized and perfused with ice-cold cutting solution (in mM) 87 NaCl, 25 NaHO_3_, 2.5 KCl, 1.25 NaH_2_PO_4_, 7 MgCl_2_, 0.5 CaCl_2_, 10 glucose, and 7 sucrose. Thereafter, the brain was immediately removed by dissection. Brain slices (300 μm) were sectioned in the coronal plane using a vibrating blade microtome (Leica Biosystems, VT1200S) in the same solution. Slices were transferred to an incubation chamber and maintained at 34 °C for 30 min and then at room temperature (23-25 °C). During whole-cell recordings, slices were continuously perfused with (in mM) 128 NaCl, 26.2 NaHO_3_, 2.5 KCl, 1 NaH_2_PO_4_, 1.5 CaCl_2_, 1.5 MgCl_2,_ and 11 glucose, maintained at 30.0±0.5°C. All solutions were equilibrated and maintained with carbogen gas (95% O_2_/5% CO_2_) throughout.

### Electrophysiology

Pyramidal neurons were targeted for somatic whole-cell recording in the CA3c region of hippocampus using gradient-contrast video-microscopy on custom-built or commercial (Bruker) upright microscopes. This region was selected based on high level of Camk2a expression in these neurons^61^. Electrophysiological recordings were obtained using Multiclamp 700B amplifiers (Molecular Devices). Signals were filtered at 10 kHz and sampled at 50 kHz using a Digidata 1440B (Molecular Devices) digitizer. For whole-cell recordings, borosilicate patch pipettes were filled with an intracellular solution containing (in mM) 124 potassium gluconate, 2 KCl, 9 HEPES, 4 MgCl_2_, 4 NaATP, 3 L-Ascorbic Acid, and 0.5 NaGTP. Pipette capacitance was neutralized in all recordings and electrode series resistance compensated using bridge balance in current-clamp mode. Recordings with series resistance > 20MΩ were discontinued. Constant current injection maintained the membrane potential of pyramidal neurons during whole cell recording at ∼-65 mV. Action potentials trains were initiated by somatic current injection (300 ms). To measure passive parameters in each recording, a -20pA current step was utilized. For analysis of all passive and active parameters, Clampfit software (Molecular Devices) and custom python scripts were utilized.

## Funding

Research reported in this publication was supported by the National Institute on Aging of the National Institutes of Health and National Institutes of Health: F32AG064862 (S.Rayaprolu), R01 NS114130 (SR), K08 NS099474 (SR), RF1 AG071587 (SR, NTS), R01AG061800 (NTS), RF1AG062181 (NTS), U01AG061357 (AIL, NTS), U54AG065187 (PI: AIL; RB), 1 R01 1R01NS115994 (LBW), NSF CAREER 1944053 (LBW), 1 R56 AG072473-01 (MR); Goizueta Alzheimer’s Disease Research Center: P30 AG066511 (PI: AIL; RB), 00100569 (MR); P50 AG025688 (PI: AIL - Pilot award to SR and LBW).

## Author Contributions

Conceptualization: S.Rayaprolu, NTS, S.Rangaraju

Methodology: S.Rayaprolu, SB, RB, SS, LH, HX, PB, DMD, RN, AMG, VJO, NTS, S.Rangaraju

Investigation: S.Rayaprolu, SB, RB, SS, LH, HX, PB, DMD, RN, AMG, VJO, NTS, S.Rangaraju

Writing-Original draft: S.Rayaprolu, NTS, S.Rangaraju

Writing-Review and Editing: S.Rayaprolu, NTS, S.Rangaraju, SS, RB, SB, PB, DMD, MR, AIL, LBW

Funding acquisition: S.Rayaprolu, S.Rangaraju, NTS, AIL, LBW

Resources: S.Rangaraju, NTS, AIL, LBW, MR

Supervision: S.Rangaraju, NTS

## Acknowledgements

Juliet Santiago and Karolina Piotrowska-Nitsche, PhD. Research reported in this publication was also supported in part by the Emory Integrated Proteomics Core (EIPC), Emory Integrate Genomic Core (EIGC), Mouse Transgenic and Gene Targeting Core (TMF), and Emory University Integrated Cellular Imaging (ICI) Microscopy Core which are subsidized by the Emory University School of Medicine and is one of the Emory Integrated Core Facilities. Additional support was provided by the Georgia Clinical & Translational Science Alliance of the National Institutes of Health under Award Number UL1TR002378 (EIPC & EIGC). Additional support to TMF was provided by the National Center for Advancing Translational Sciences of the National Institutes of Health under Award Number UL1TR000454.

## Ethics approval and consent to participate

Approval from the Emory University Institutional Animal Care and Use Committee was obtained prior to all animal-related studies (IACUC protocols # PROTO201800252 and PROTO201700821).

## Consent for publication

All authors have approved of the contents of this manuscript and provided consent for publication.

## Availability of data and materials

The mass spectrometry proteomics data have been deposited to the ProteomeXchange Consortium via the PRIDE partner repository with the dataset identifier: px-submission #519326.

## Competing interests

The authors declare that they have no competing interests.

## SUPPLEMENTAL FIGURE LEGENDS

**Supplemental Figure S1.**
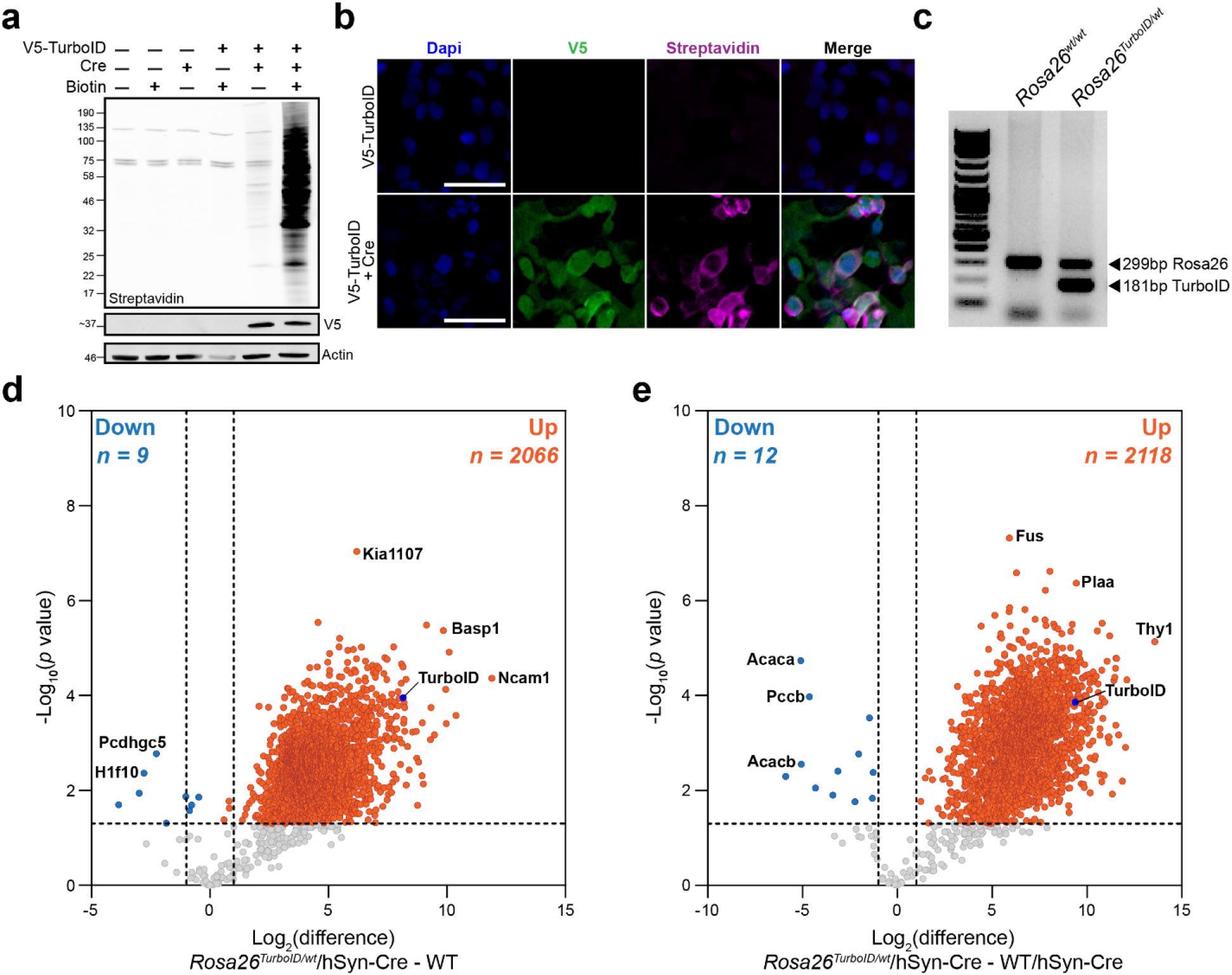
*In vitro* validation of Rosa26 TurboID targeting vector. These data associated with Figure 1. **a** Western blot studies of whole cell HEK293 lysates after transfection with Rosa26 TurboID targeting vector or sham, and co-transfection with CMV-Cre plasmid followed by biotin supplementation in culture media. Biotinylation (streptavidin680), V5 (to detect TurboID), and actin were measured. As compared to few endogenous biotinylated proteins in un-transfected or single plasmid transfected cells, only co-transfected cells treated with biotin showed biotinylation while low level biotinylation was also observed in biotin non-supplemented wells due to the presence of biotin in cell culture media. **b** Representative immunofluorescence images of cells adhered to coverslips, from experiments in (a), confirming expression of V5-TurboID-NES (green) and biotinylation (magenta) in co-transfected HEK293 cells. Scale bar = 200 µm **c** Representative genotyping PCR results obtained from *Rosa26^wt/wt^* and *Rosa26^TurboID/wt^* littermate mice. The 181bp band corresponds TurboID transgene and 299bp band represents endogenous *ROSA26* allele. **d, e** Volcano plots showing differentially expressed proteins comparing (d) *Rosa26^TurboID/wt^*/hSyn-Cre with un-injected WT mice, and (e) *Rosa26^TurboID/wt^* /hSyn-Cre mice with WT/hSyn-Cre mice. Orange symbols (*p* ≤ 0.05 and ≥ 2-fold change) represent biotinylated proteins enriched in the *Rosa26^TurboID/wt^*/hSyn-Cr e brains while blue symbols represent biotinylated proteins enriched in control groups.

**Supplemental Figure 2.**
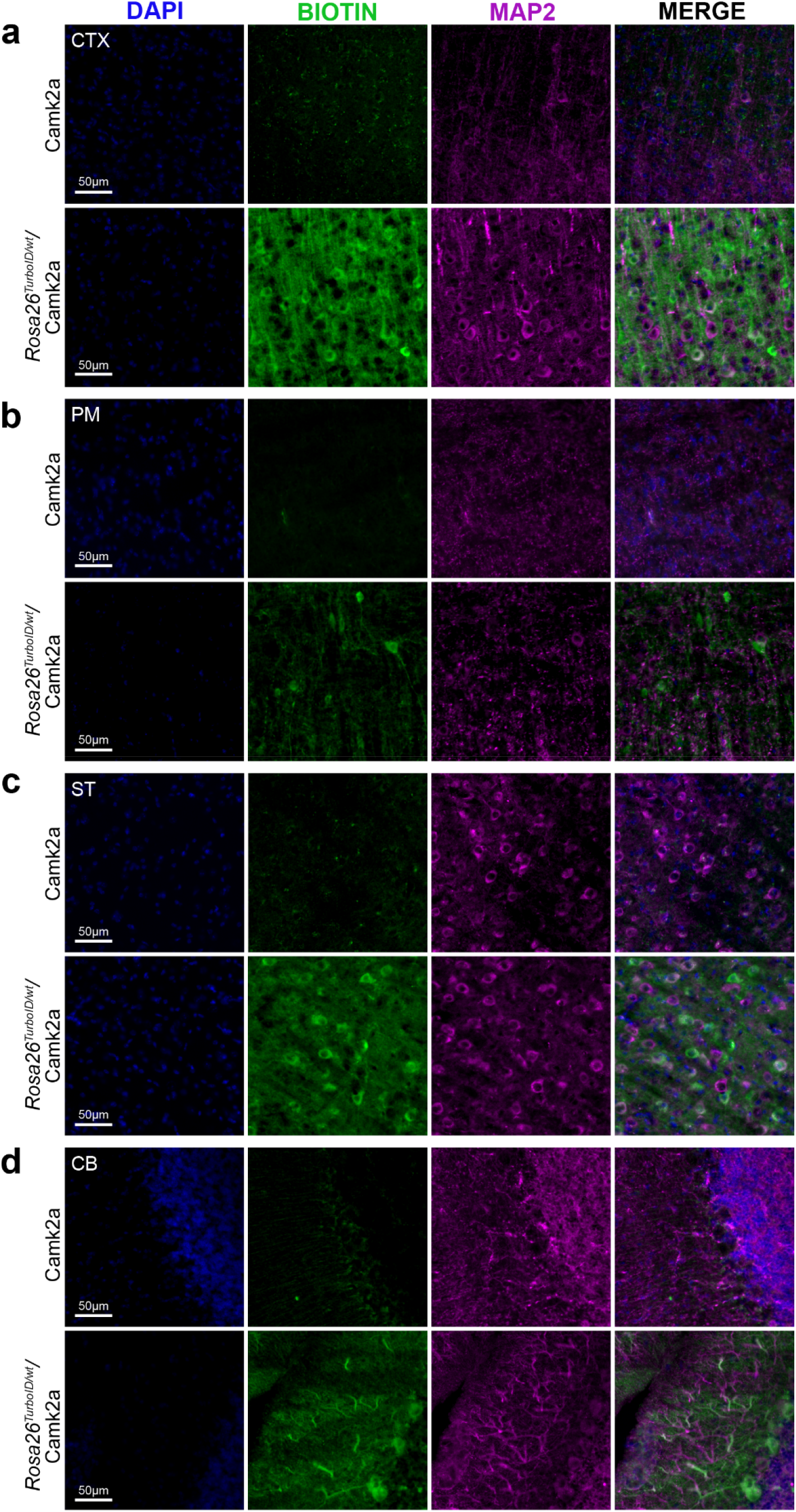
Immuno-histochemical confirmation of neuronal biotinylation in *Rosa26^TurboID/wt^*/Camk2a mice. These data are associated with Figures 2 and 3. Representative immunofluorescence images displaying robust biotinylation (green: streptavidin Alexa488) within neuronal cell bodies and axons (magenta: Map2) in the **a** cortex (CTX), **b** pons**/**medulla (PM), **c** striatum/thalamus (ST), and **d** cerebellum (CB). Degree of biotinylation varied by region, with highest biotinylation observed in the CTX and PM regions. Neurons labeled in the CTX were mostly pyramidal neurons while Purkinje neurons were labeled in the cerebellum. PM showed labeling predominantly of axons. Nuclei were labeled with DAPI (blue).

**Supplemental Figure 3.**
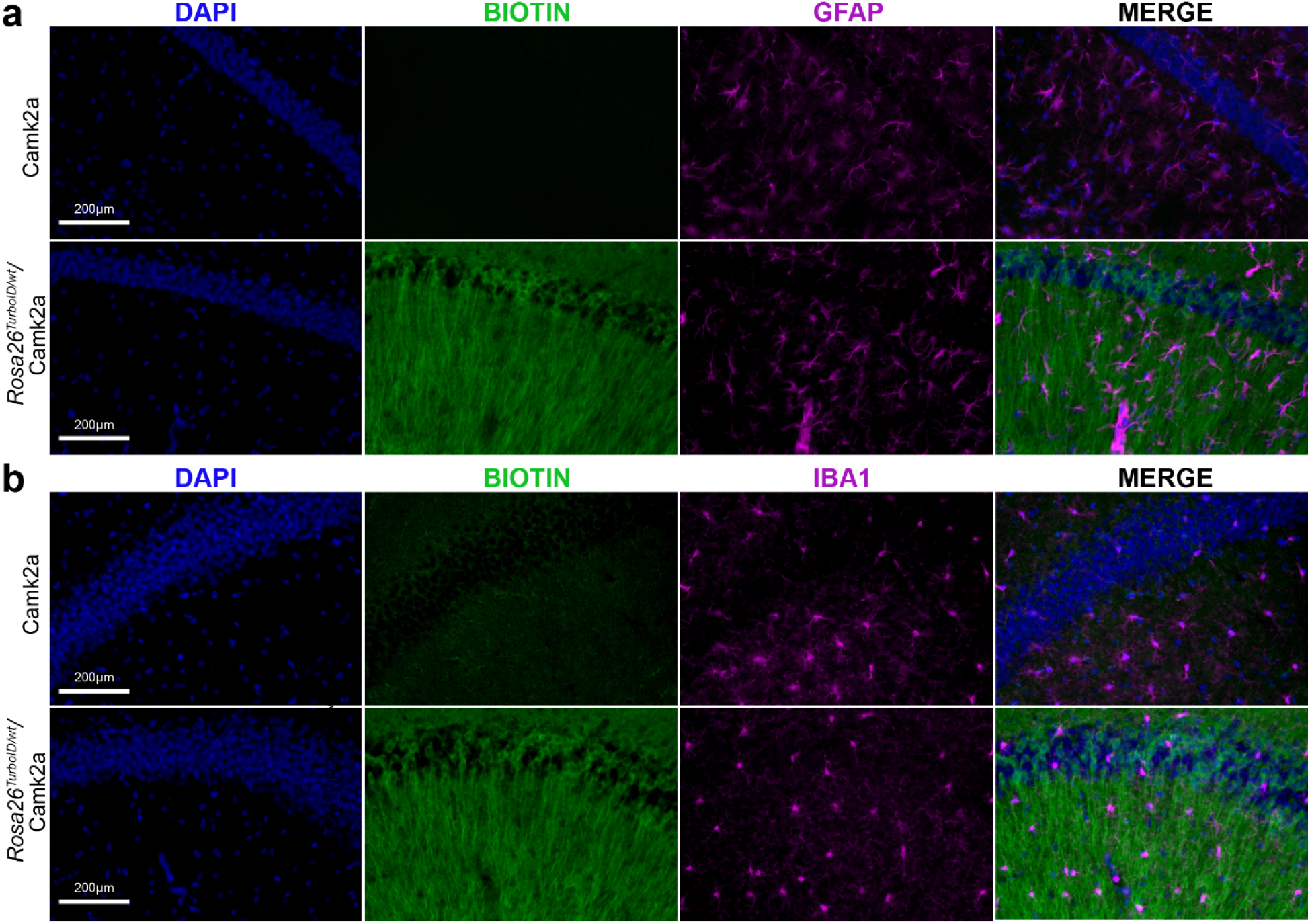
Lack of glial cell biotinyation or reactive glial morphological changes in *Rosa26^TurboID/wt^*/Camk2a mice. These data are associated with Figures 2 and 3. **a, b** Representative immunofluorescence images of CA2 of the hippocampus confirming lack of overlap between biotinylation (green: streptavidin Alexa488) with **a** astrocytes (magenta: Gfap) or **b** microglia (magenta: Iba1). Nuclei were labeled with DAPI (blue).

**Supplemental Figure 4.**
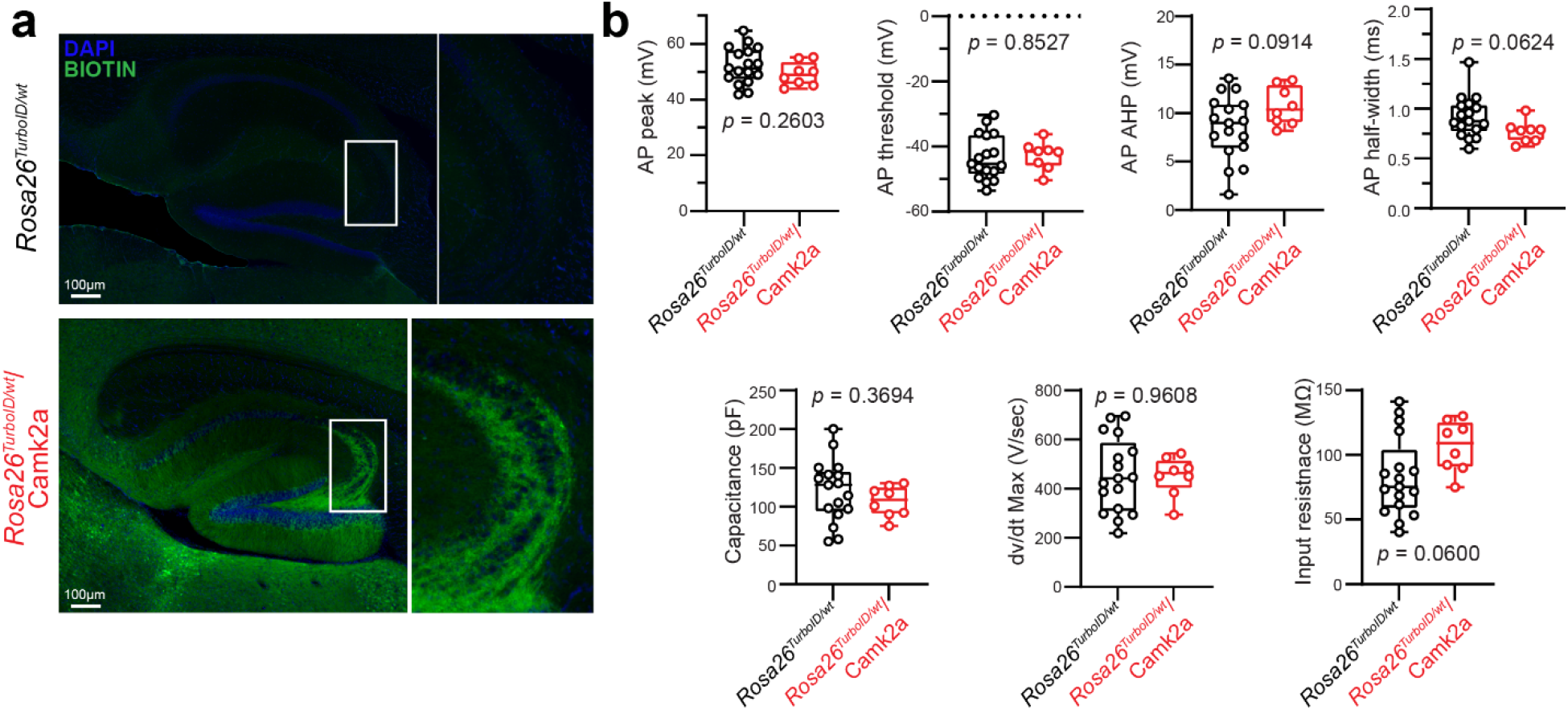
Lack of electrophysiological alterations in hippocampal CA3 pyramidal neurons in *Rosa26^TurboID/wt^*/Camk2a mice. These data are associated with Figure 2. **a** Representative immunofluorescence images of the hippocampus from control (*Rosa^TurboID/wt^*) and *Rosa26^TurboID/wt^*/Camk2a-Cre^Ert2^ displaying biotinylation within cells, specifically CA3 neurons. **b** Summary data from whole-cell current clamp recordings in CA3c in control (*Rosa^TurboID/wt^*) and labeled *Rosa26^TurboID/wt^*/Camk2a-Cre^Ert2^ mice. Each data point represents a single neuron. Pooled analysis from *n* = 17 non-labeled control and *n* = 8 labeled neurons (*n* = 2 mice/group; *p* value in each graph represents unpaired t-test).

**Supplemental Figure 5.**
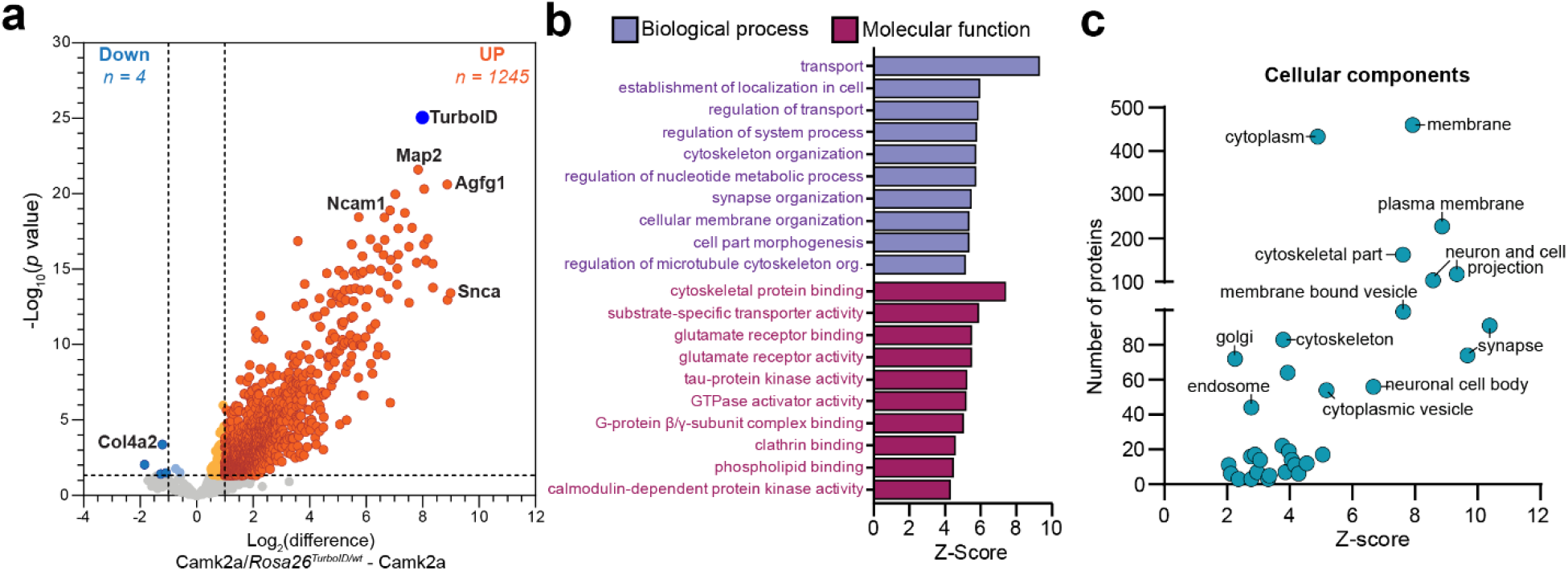
*In vivo* Camk2a-positive neuronal proteomic biotinylation by TurboID provides a representative cellular proteome. **a** Volcano plot of MS data showing differentially enriched proteins between labeled *Rosa26^TurboID/wt^*/Camk2a-Cre^Ert2^ and non-labeled Camk2a-Cre^Ert2^ control mouse brain. For this analysis, all 5 brain regions were combined for both groups. Orange symbols (*p* ≤ 0.05 and ≥ 2-fold change) represent biotinylated proteins enriched in the *Rosa26^TurboID/wt^*/Camk2a-Cre^Ert2^ brain while blue symbols represent biotinylated proteins enriched in control brain. **b** GSEA of biotinylated proteins (orange symbols in panel a) showed neuron-specific gene ontology terms as well as metabolic, cytoskeletal, ion transporter, and endocytosis related gene ontology terms. **c** Cellular component gene ontology terms enriched in Camk2a proteome showing several cellular sub-compartments and organelles as well as the synapse. For related MS data and additional analyses, see Supplemental Tables 10 & 11.

**Supplemental Figure 6.**
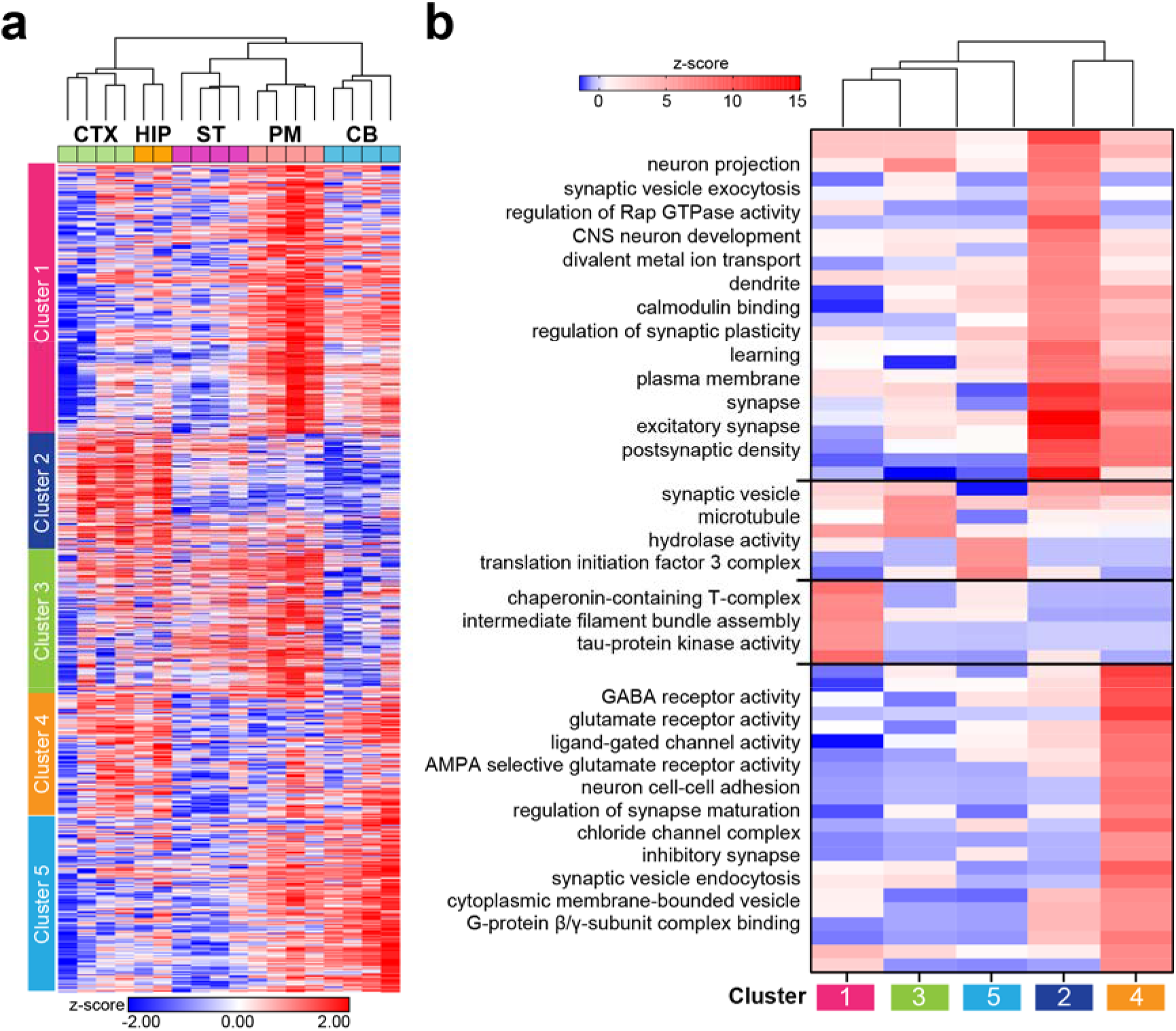
Region-specific proteomic differences in adult mouse Camk2a. **a** Unsupervised hierarchical clustering analysis (HCA) of 1,245 Camk2a neuronal proteins (based on rows and columns) from *Rosa26^TurboID/wt^*/Camk2a-Cre^Ert2^ mice showed distinct clusters of proteins, primarily highlighting region-specific patterns of protein expression. Prior to HCA, MS data was normalized to TurboID abundance per sample to account for inherent differences in TurboID expression and/or Camk2a promoter activity across regions. Individual clusters are shown with distinct colors. **b** Heat map representation (HCA) of results from GSEA to identify over-represented molecular functions, cellular components, and biological processes within clusters shown in panel (a). Representative gene ontology terms are highlighted. For related MS data and additional analyses, see Supplemental Tables 17 & 18.

## Notes

### Competing Interest Statement

The authors have declared no competing interest.

